# Amplified cortical tracking of word-level features of continuous competing speech in older adults

**DOI:** 10.1101/2020.12.21.423513

**Authors:** Juraj Mesik, Lucia A. Ray, Magdalena Wojtczak

## Abstract

Speech-in-noise comprehension difficulties are common among the elderly population, yet traditional objective measures of speech perception are largely insensitive to this deficit, particularly in the absence of clinical hearing loss. In recent years, a growing body of research in young normal-hearing adults has demonstrated that high-level features related to speech semantics and lexical predictability elicit strong centro-parietal negativity in the EEG signal around 400 ms following the word onset. Here we investigate effects of age on cortical tracking of these word-level features within a two-talker speech mixture, and their relationship with self-reported difficulties with speech-in-noise understanding. While undergoing EEG recordings, younger and older adult participants listened to a continuous narrative story in the presence of a distractor story. We then utilized forward encoding models to estimate cortical tracking of three speech features: 1) “semantic” dissimilarity of each word relative to the preceding context, 2) lexical surprisal for each word, and 3) overall word audibility. Our results revealed robust tracking of all three features for attended speech, with surprisal and word audibility showing significantly stronger contributions to neural activity than dissimilarity. Additionally, older adults exhibited significantly stronger tracking of surprisal and audibility than younger adults, especially over frontal electrode sites, potentially reflecting increased listening effort. Finally, neuro-behavioral analyses revealed trends of a negative relationship between subjective speech-in-noise perception difficulties and the model goodness-of-fit for attended speech, as well as a positive relationship between task performance and the goodness-of-fit, indicating behavioral relevance of these measures. Together, our results demonstrate the utility of modeling cortical responses to multi-talker speech using complex, word-level features and the potential for their use to study changes in speech processing due to aging and hearing loss.

## 1. Introduction

Speech perception is fundamentally important for human communication. While speech signals are often embedded in complex sound mixtures that can interfere with speech perception via energetic and informational masking, the auditory system is remarkably adept at utilizing attentional mechanisms to suppress distractor information and enhance representations of the target speech (e.g., Ding and Simon, 2012a; Mesgarani and Chang, 2012; O’Sullivan et al., 2019). However, the robustness of speech perception, particularly in the presence of noise, is vulnerable to deterioration through both noise-induced and age-related hearing loss (Dubno et al., 1984; Helfer and Wilber, 1990; Fogerty et al., 2015, 2020) as well as age-related cognitive decline (van Rooij and Plomp, 1990; Akeroyd, 2008; Dryden et al., 2017). Additionally, a small but significant portion of the population experiences speech-in-noise (SIN) perception difficulties, without exhibiting clinical hearing loss (Saunders, 1989; Zhao and Stephens, 2007; Tremblay et al., 2015). Together, these SIN perception difficulties can lead to significant impairment in quality of life (Dalton et al., 2003; Chia et al., 2007), and in older adults they may result in increased social isolation (Chia et al., 2007; Mick et al., 2014; Pronk et al., 2014), potentially exacerbating loss of cognitive function (Loughrey et al., 2018; Ray et al., 2018).

Although subjective SIN perception difficulties are relatively common in older individuals, objective tests for quantifying these deficits, such as identification of words or sentences in noise (e.g., QuickSin; Killion et al., 2004), often do not strongly correlate with the degree of subjective deficit (Phatak et al., 2018), particularly in cases with little-to-no clinical hearing loss. Smith and colleagues (2019) recently reported that only 8% of their sample of 194 listeners exhibited deficits in objective SIN tasks, while 42% of listeners indicated experiencing subjective SIN perception difficulties. A likely reason for this mismatch is that objective speech perception tests do not accurately reflect real world scenarios where SIN difficulties arise. For example, while existing tests generally require identification of isolated words or sentences embedded in noise (e.g., speech-shaped noise or a competing talker), real world speech perception often requires real-time comprehension of multi-sentence expressions, embedded in a reverberant environment, in the presence of multiple competing speakers at different spatial positions. In these scenarios, listeners who need to expend additional time and cognitive resources to identify the meaning of the incoming speech may “fall behind” in comprehension of later parts of the utterance. Moreover, even if the listener can correctly piece together the meaning of the utterance, their subjective confidence may be diminished, potentially “blurring” the predictive processes thought to facilitate perception of upcoming speech (Pickering and Gambi, 2018). As such, behavioral measures that more accurately reflect subjective SIN perception difficulties may require utilization of more realistic, narrative stimuli, and focus on quantifying comprehension, as opposed to simple word or sentence identification (e.g., Xia et al., 2017).

While development of behavioral paradigms focusing on characterizing SIN perception difficulties is an important goal, a complementary and potentially more sensitive approach to quantifying these deficits may be provided by neural measures of continuous-speech tracking. In recent years, non-invasive methodologies for measurement of neural representations of continuous speech in humans have become increasingly popular (Lalor and Foxe, 2010; Crosse et al., 2016), particularly in application to young normal-hearing (YNH) populations. One important result of this work has been the demonstration of profound attentional modulation of speech whereby temporal dynamics of neural responses to attended and ignored speech differ considerably, both in representation of lower-level features such as the speech envelope (Ding and Simon, 2012; Power et al., 2012; Kong et al., 2014; Fiedler et al., 2019), and higher-level features related to lexical and semantic content of speech (Brodbeck et al., 2018; Broderick et al., 2018). Indeed, while lower-level features produce robust responses even when speech is ignored, features related to linguistic representations only show robust responses for attended speech, suggesting that they are tightly linked with speech comprehension. Responses to higher-level features may therefore be particularly sensitive to SIN perception difficulties, which are likely associated with impaired comprehension performance. In fact, SIN perception difficulties could potentially manifest themselves not only in terms of poorer tracking of higher-level features in attended speech, but also in increased tracking of features in ignored speech, when facing difficulties with suppression of distractor information.

Changes in neural processing of continuous speech in aging populations, compared to young adults, are relatively poorly understood. Several studies have utilized magneto- and electroencephalography (M/EEG) to address this question. Studies comparing envelope-related cortical responses have revealed a pattern of amplified envelope representations in older populations (Presacco et al., 2016; Decruy et al., 2019; Zan et al., 2020), potentially reflecting changes in the utilization of cognitive resources during speech comprehension. More recently, Broderick et al. (2020) compared higher-level representations of speech in younger and older populations. They estimated EEG responses to 5-gram surprisal, reflecting the predictability of words given the preceding sequence of four words, as well as semantic dissimilarity, reflecting the contribution of each word to the semantic content of a sentence. While younger listeners showed strong responses to both of these features, older adults exhibited a delayed surprisal response and a near-absent response to semantic dissimilarity. These findings demonstrate that representations of higher-level features of speech may indeed reveal robust effects of age. However, because Broderick et al. (2020) did not report behavioral measures related to speech comprehension, nor measures of subjective speech perception difficulties among their participants, it is unclear whether these metrics would correlate with the reported EEG-based findings. Moreover, participants in that study were presented with clear speech without any distractors (e.g., competing speakers), making it unclear how speech representations differ in complex listening scenarios where speech perception difficulties are most commonly reported.

The goal of this study was to compare higher-level neural representations of two-talker speech mixtures between younger and older adults, and to explore how these measures relate to comprehension performance and self-reported SIN perception difficulties. In particular, we examined representations related to word dissimilarity relative to short-term preceding context, lexical surprisal based on multi-sentence context, and word-level audibility. We chose to pursue this paradigm for several reasons. First, a multi-talker paradigm was chosen because subjective SIN perception difficulties commonly arise in aging listeners in the context of competing speech. If age-related changes in neural representations are confirmed, then these neural signatures could potentially be further explored as a candidate objective correlate for subjective SIN difficulties. Second, we chose to characterize responses to word-level features linked to meaning and lexical predictability because existing evidence indicates that responses to higher-level features are tightly linked to speech comprehension (Broderick et al., 2018). As such, we anticipated that responses to these features are more likely to exhibit differences as a function of age and SIN perception difficulties. Although neural representations reflecting the end-goal of speech perception may allow for only limited inference about the underlying causes of SIN perception difficulties, which can range from peripheral changes in acoustic representations to more central changes in cognitive processes, these representations may offer increased sensitivity due to capturing the combined effects of the various etiologies underlying the deficit.

## 2. Materials and Methods

### 2.1 Participants

In total, 45 adult volunteers completed the experiment, and data from 41 participants were used due to a methodological change implemented early in data collection. The participant pool was divided into two groups, younger adults (YA) and older adults (OA), with participants who were 18-39 years included in the former, and participants who were 40-70 years included in the latter. The YA group consisted of 20 participants (6 male, 14 female; mean ± s.d. age: 29.40 ± 6.40 years), while the OA group included 21 participants (9 male, 12 female; mean ± s.d. age: 53.48 ± 8.68 years). Participants were recruited via email advertisement from a pool of students, staff, and alumni of the University of Minnesota. All participants provided informed written consent and received either course credit or monetary compensation for their participation. The procedures were approved by the Institutional Review Board of the University of Minnesota.

### 2.2 Audiometry

An air-conduction audiogram was measured in each ear for each participant prior to beginning the EEG procedures. Detection thresholds were measured at octave frequencies in the 250 – 8000 Hz range, and frequencies for which thresholds exceeded 20 dB HL were deemed to be affected by hearing loss (HL). This procedure resulted in the detection of 2 participants in the YA group, and 16 participants in the OA group as having mild-to-moderate high-frequency HL. The skewed distribution of HL towards the older population was expected, as peripheral frequency sensitivity naturally diminishes with age (see reviews by Huang and Tang, 2010; Yamasoba et al., 2013).

For participants with any hearing loss, all experimental audio materials were amplified in the frequency regions of hearing loss, as described in section 2.4 below. Under these conditions, we observed no association between task performance and high-frequency hearing loss.

### 2.3 Modified SSQ questionnaire

Prior to the EEG procedures, all participants completed a modified version of a subset of Speech, Spatial and Qualities of Hearing Scale (SSQ_m_). The original version of SSQ (Gatehouse and Noble, 2004) was designed to measure subjective hearing challenges faced by listeners in various situations of daily life. In our version, we specifically probed participants about difficulties with and frustrations related to hearing speech in noisy situations, such as cafes and social gatherings. Each of the 14 items was presented on a computer screen along with four graded choices of frequency, difficulty, or discomfort related to the presented listening scenarios. E.g.,

#### Item 1

I find it difficult to talk with staff in places such as shops, cafes, or banks, due to struggling to hear what they are saying.

#### Item 10

In group conversations I worry about mishearing people and responding based on incorrect information.

#### Response choices

1. Not at all
2. Rarely
3. Often
4. Very often

### 2.4 Stimuli

Stimuli were four public domain short story audiobooks (*Summer Snow Storm* by Adam Chase; *Mr. Tilly’s Seance* by Edward F. Benson; *A Pail of Air* by Fritz Leiber; *Home Is Where You Left It* by Adam Chase; source: LibriVox.org), spoken by two male speakers (two stories per speaker). Each story was about 25 min in duration and was pre-processed to truncate any silences between words that exceeded a 500-ms interval to 500 ms. On a block-by-block basis (see section 2.5 below), each audiobook was root-mean-square (RMS) normalized and scaled to 65 dB SPL. Stimuli were presented to participants using ER1 Insert Earphones (Etymotic Research, Elk Grove Village, IL), shielded with copper foil to prevent electrical artifacts in the EEG data.

In order to minimize the odds of finding age-related differences in neural responses that could be attributed to reduced audibility in participants with hearing loss, all audio materials were custom-filtered for each participant with HL using a FIR filter implemented in MATLAB (Mathworks, Natick, MA) via the *designfilt* and *filter* functions. The filter was designed to apply half gain, amplifying all frequency bands by half the amount of the hearing loss:

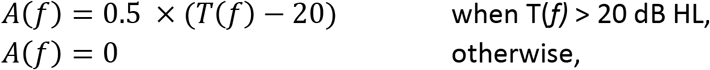

where *T*(*f*) is the detection threshold in dB HL at frequency *f*. Note that half gain amplification is a commonly used strategy to mitigate reduced audibility due to hearing loss, while preventing discomfort from loudness recruitment, whereby loudness growth for frequencies affected by cochlear hearing loss is steeper than that observed in normal hearing (Fowler, 1936; Steinberg and Gardner, 1937).

### 2.5 Experimental procedures

The experimental setup was implemented using the Psychophysics Toolbox (Brainard, 1997; Pelli, 1997; Kleiner et al., 2007) in MATLAB. Two experimental runs were completed by each study participant. In each run, a pair of audiobooks read by different male speakers (Fig. 1A) was presented diotically (the mixture of the two audiobooks in each ear) to the participant. One of the stories served as the *attended* story, while the other was the *ignored* story, with these designations being counter-balanced across participants. A run was broken up into 24-27 blocks (variation was due to small differences in durations of audiobooks used in each of the two runs). Each block contained a roughly 1-minute segment of audio, followed by a series of questions, detailed below. Block duration was allowed to exceed 1 minute in order to ensure that each block concluded at the end of a sentence in the attended story. The attended story remained the same throughout the run. To cue the participants to follow the correct story, the audio of the attended story started 1 sec prior to the onset of the ignored story. This was further aided by making this initial 1-sec portion of the attended story in each block (except block #1) correspond to the final 1-sec of the attended story from the previous block. These repeated segments with the attended story alone were excluded from statistical analyses. Throughout each block, participants were instructed to stay as still as possible, and to keep their gaze on a central fixation marker presented on a computer display in front of the participant. The purpose of this was to minimize EEG artifacts caused by muscle activity.

**Figure 1.**
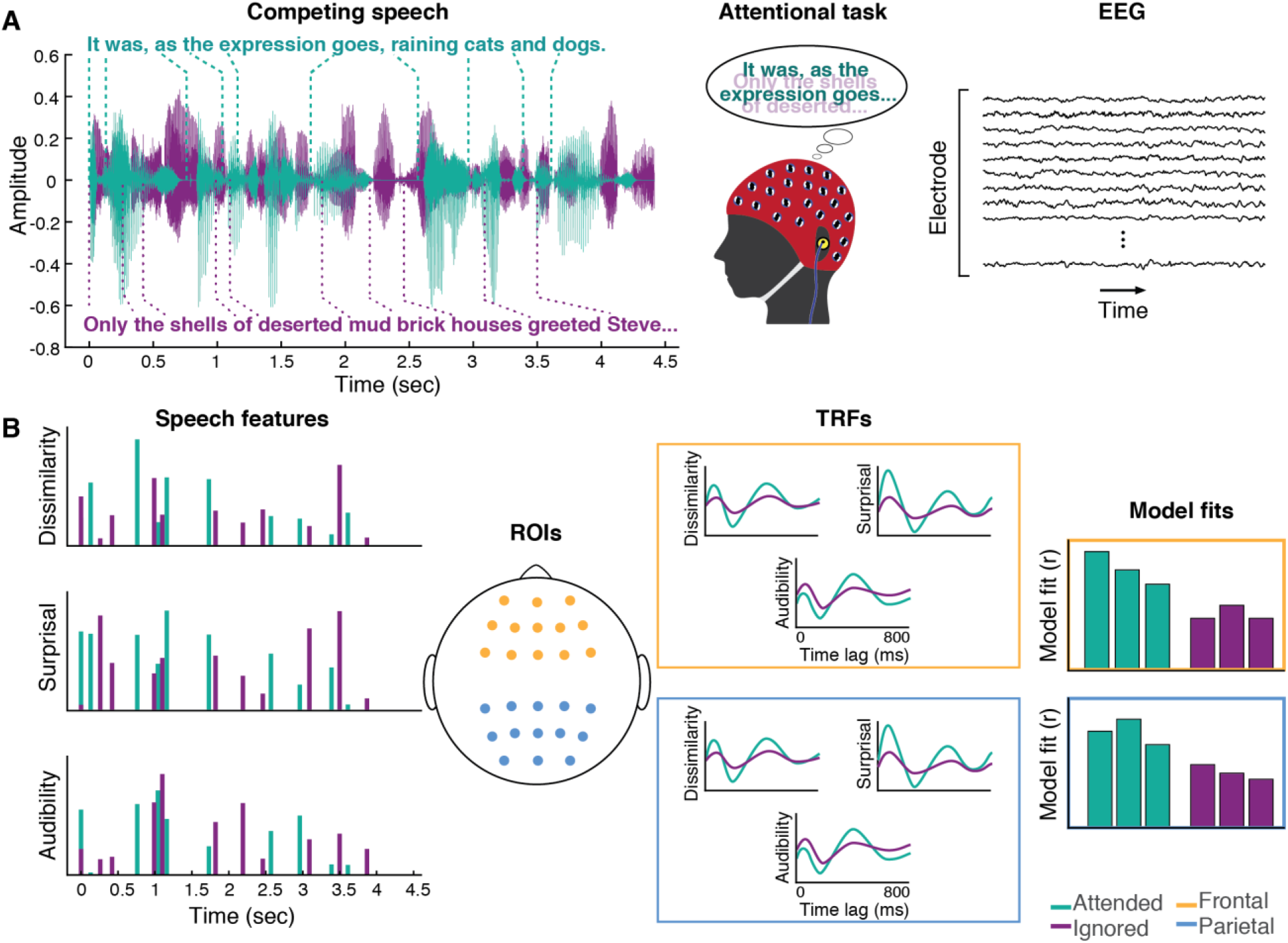
Experimental procedures. (A) Participants listened to a mixture of two speakers, while attending to one of them. Meanwhile, 64-channel EEG was recorded from their scalp. (B) Three word-level features (dissimilarity, surprisal, and audibility) were extracted from the speech for both the attended and ignored stories, and used to generate regressors containing impulses that were time-aligned to the word onsets scaled by the amplitude of each feature. These features were regressed against the EEG signals recorded during the experiment, resulting in TRF and model fit contributions for each of the features. These TRFs and goodness-of-fit values were averaged across groups of frontal (yellow) and parietal (blue) electrodes for use in group-level analyses.

Following each block, participants were presented on a display with a series of Yes/No questions about the audio from that block, including:

1. Four comprehension questions about the contents of the attended story
2. Confidence ratings for each of the comprehension questions
3. Intelligibility judgment about the attended speaker
4. Subjective attentiveness rating

As each behavioral question had binary answer choices (e.g., for attentiveness, participants answered “Were you able to stay focused on the target story?” Yes/No), the main purpose of these questions was to gather information about participants’ comprehension and subjective experience throughout the run, and to make sure that they were attending to the correct story.

Participants were given 10 seconds to answer each question using a key press. If 10 seconds elapsed without a response, the question was marked as no-response. After answering each block’s questions, participants were allowed to request a short break to ensure that they remained comfortable throughout the experiment. These breaks were limited to up to two minutes, during which participants remained seated. The next block started as soon as the break was terminated by the participant with a key press, or two minutes elapsed. Furthermore, between the two experimental runs, participants were offered an extended break inside the booth. The EEG cap and the insert phones were not removed during the breaks.

The second experimental run was procedurally identical to the first one, except a different pair of stories was presented, neither of which was used in the first run. Additionally, the attended and ignored speakers were switched, so that the speaker that narrated the ignored story in the first run was attended in the second run, while the attended speaker from the first run became the ignored speaker in the second run. Participants were explicitly informed of this switch, and the purpose of this was to balance any possible speaker effects on each participant’s EEG data.

### 2.6 EEG procedures

While engaging in the experimental task described above, each participant’s EEG activity was sampled at 4096 Hz from their scalp using a Biosemi ActiveTwo system (BioSemi B.V., Amsterdam, The Netherlands), with 64 channels positioned according to the international 10-20 system (Klem et al., 1999). Additional external electrodes were placed on the left and right mastoids, and above and below the right eye (vertical electro-oculogram, VEOG). Prior to the beginning of the recording, and between the two runs, the experimenter visually inspected signals in all electrodes, and for any electrodes with DC offsets exceeding ± 20 mV, the contact between the electrode and scalp was readjusted until the offset fell below ± 20 mV.

### 2.7 EEG preprocessing

All pre-processing analyses were implemented via the EEGLAB toolbox (Delorme and Makeig, 2004) for MATLAB, unless otherwise stated. To reduce computational load, the raw EEG data were initially downsampled to 256 Hz, and band-pass filtered between 1 and 80 Hz using a Hamming windowed sinc FIR filter implemented in the *pop_eegfiltnew* function of EEGLAB. Subsequently, data were pre-processed using the PREP pipeline (Bigdely-Shamlo et al., 2015). These steps included line noise removal, detection of disproportionately noisy channels via an iterative robust referencing procedure, interpolation of noisy channels, and referencing the data using the final “clean” estimate of the global mean activation. The benefit of this procedure is that it minimizes the risk of signal contamination from electrodes with abnormal signals (e.g., due to faulty hardware) during the referencing stage.

Next, activations from all experimental blocks were epoched and independent component analysis (ICA; Jutten and Herault, 1991; Comon, 1994) was applied to the data using the infomax ICA algorithm (Bell and Sejnowski, 1995) implementation in EEGLAB. This procedure decomposes the EEG signal into statistically independent sources of activation, some of which reflect sensory and cognitive processes, while others capture muscle-related signal contributions and other sources of noise. We removed all components that matched eye-blink related activity in component topography, amplitude, and temporal characteristics, as well as other high-amplitude artifacts that reflected muscle activity. This, on average, led to the removal of 2.52 (SD: 0.97) components.

The cleaned EEG signals were then band-pass filtered between 1 and 8 Hz with a Chebyshev type 2 filter designed using MATLAB’s *designfilt* function (optimized to achieve 80 dB attenuation below 0.5 Hz and above 9 Hz, with pass-band ripple of 1 dB), and applied to the data using the *filtfilt* function. Afterwards, the data were z-scored in order to control for inter-subject variability in the overall signal amplitude due to nuisance factors such as skull thickness or scalp conductivity, as well as to improve efficiency in the cross-validated regression and ridge parameter search for deriving the temporal response function (TRF), described below (section 2.9.1). Finally, because run duration varied slightly due to unequal lengths of the two pairs of audiobooks (i.e. 24-27 minutes), in order to equalize contributions from each run to the overall analysis results, only blocks 2-23 from each run were used in the remaining analyses. The first block was excluded in order to minimize effects of initial errors in attending to the target story, which happened to a very small number of participants (less than 5), but was quickly corrected after initial comprehension questions were presented.

### 2.8 Word timing estimation

Word onset timings for all words within each story were estimated using the Montreal Forced Aligner (McAuliffe et al., 2017). Prior to running the aligner, the audiobook text was preprocessed to remove punctuation, typographic errors and abbreviations, and both the text and audio were divided into roughly 30-sec segments. This segmented alignment approach was used in order to prevent accumulation of alignment errors for later portions of the audio. All alignments were subsequently manually inspected for timing errors, and when noticeable alignment errors were detected, the aligner was re-run on further-shortened (15 sec) segments of the affected audio. While forced alignment routinely results in some degree of timing errors, these are typically small, with a median of about 15 ms for the aligner used here. As such, only a small degree of temporal smearing of estimated neural responses should occur due to these errors.

### 2.9 Data analysis

#### 2.9.1 TRF analyses

Time courses of cortical responses to different speech features, known as the TRFs, were extracted from preprocessed EEG activity using cross-validated regularized linear regression, implemented via the mTRF toolbox (Crosse et al., 2016). Briefly, deconvolution of a TRF for a given feature from the EEG signal is accomplished by first constructing a regressor containing a time series, sampled at a rate matching the EEG signal, of that feature’s amplitudes. By including multiple time-lagged copies of the regressor for each feature, the effect of a given feature on the neural activity at different latencies relative to the word onset can be estimated, resulting in a time course of neural response. Regressors for all features are combined into a full design matrix, and this matrix is then regressed against the EEG signal to yield the impulse responses (i.e., TRFs) for each of the included features at each electrode site.

In practice, this procedure was implemented through 11-fold cross-validation, with each fold involving three steps. First, the data and regressors were split into a training set, composed of 40 blocks of the data (~40 minutes), and a testing set, containing the remaining 4 blocks of the data (~4 minutes). Next, the training set was used to determine the ridge parameter, λ, by iteratively fitting the cortical-response model using a range of ridge parameters. The TRF estimates were obtained for the λ parameter that produced the best model fit to the training data, as determined by the highest Pearson’s correlation coefficient between the predicted and actual EEG signal. The TRF estimates were then used to assess the model fit for the test data. This was done by convolving the estimated TRFs with the corresponding word-feature regressors for the test data set, and computing the Pearson’s correlation between the predicted and actual test data. Following cross-validation, average TRFs for each feature and an average model goodness-of-fit were computed from results of all cross-validation folds for use in group-level analyses.

##### 2.9.1.1 Regression features

Word features used in the regression analyses included semantic dissimilarity, surprisal, and word audibility (Fig. 1B).

###### 2.9.1.1.1 Semantic dissimilarity

Semantic dissimilarity, reflecting approximately the degree to which each word adds new information to a sentence, was computed as described in Broderick et al., (2018). Briefly, we used Google’s pre-trained *word2vec* neural network (Mikolov et al., 2013a, 2013b), implemented using the Gensim library (Rehurek and Sojka, 2010) for Python, to compute a 300-dimensional vector representation (otherwise known as an embedding) of each word within our stimuli. An important property of these vector representations is that in the 300-dimensional vector space, vectors of words with similar meanings point in similar directions. Computing correlation between vectors representing any two words approximates their semantic similarity. Because EEG response to incongruent words has been shown to elicit a strong N400 component (Kutas and Hillyard, 1980), for regression purposes these similarity values were subtracted from 1 to convert them to dissimilarity.

To construct semantic dissimilarity regressors, we computed the dissimilarity between each word’s vector, and the average of vectors for all preceding words in a given sentence. In the case of the first word in a sentence, we computed dissimilarity from the average vector for words in the previous sentence. These dissimilarity values were then used to construct the regressor consisting of unit-length impulses aligned to word onsets that were scaled by each word’s dissimilarity value and zeros between these impulses. Although neural responses to semantic content of words may not be strictly time-locked to word onsets, potentially leading to some degree of temporal smearing in the estimated TRFs, word onset timings have been successfully used as timestamps for characterizing higher-order lexical and semantic processes (e.g., Broderick et al., 2018; Weissbart et al., 2019).

###### 2.9.1.1.2 Lexical surprisal

Surprisal regressors were constructed in an identical way to dissimilarity, except the feature values were computed using OpenAI’s GPT-2 (Radford et al., 2019; 12-layer, 117M parameter version) artificial neural network (ANN), similar to the approach demonstrated by Heilbron et al. (2019). These procedures were implemented in Python using the Transformers library (Wolf et al., 2020) for PyTorch (Paszke et al., 2019). GPT-2 is a transformer-based (Vaswani et al., 2017) ANN that, using a “self-attention” mechanism, is capable of effectively using hundreds of words worth of preceding context in order to generate seemingly realistic sequences of text. As a result, it can be used as a proxy for computing the predictability of words within a sequence. Surprisal is calculated based on a much longer time scale (a large number of words in the preceding context) than semantic dissimilarity. Specifically, by providing GPT-2 with a segment of text and then generating the distribution over the next word, it is possible to assess the relative probability of the actual next word within GPT-2’s distribution of possibilities. Generation of all probabilities involves iteratively adding words into the context, and computing the probability of each successive word. In practice, GPT-2 utilizes a tokenized representation of text, whereby GPT-2’s vocabulary corresponds to a combination of whole words (particularly in the case of shorter words) and word fragments.

As a result, the probability of the i-th word *w_i_* was computed as a product of conditional probabilities of the constituent word tokens *t*, with each token’s probability being computed with the model’s knowledge of the preceding tokens (i.e. preceding text plus current word’s tokens whose probabilities were already estimated):

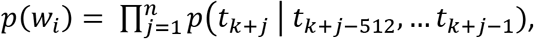

where *j* indexes the *n* tokens of word *w_i_*, *k* is the absolute index of the last token in the preceding word (relative to text beginning), and 512 is the maximum number of tokens utilized for prediction. For token indices less than 512 (i.e., early portions of the text), all of the available context was used. Furthermore, in cases where one or more tokens from the word at the far boundary of the context window did not fit into the 512 token limit, that word’s tokens were excluded from being used for prediction. Note that although GPT-2 is capable of utilizing up to 1024 tokens for prediction, we utilized a context length of 512 tokens due to limited computational resources. Across the 4 stories, when full predictive context was utilized for prediction, it contained on average 393.3 [s.d. = 31.1] words.

Because brain mechanisms underlying lexical prediction respond more to unexpected than to expected words (Kutas and Hillyard, 1984), surprisal was computed by taking the negative log of the conditional probabilities of each word, leading to less expected words receiving higher surprisal values:

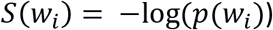

###### 2.9.1.1.3 Audibility

Word audibility regressors were constructed separately for the attended and ignored stories to capture the degree of masking of each word in one story by the speaker of the other story. In contrast to dissimilarity and surprisal, this value reflects the information at the shortest, word-by-word time scale, with higher signal-to-noise ratio (SNR) values reflecting greater peripheral fidelity of target speech, leading to lower uncertainty in speech identification on the basis of the bottom-up signal. For each word *w_i_* in a given story, its audibility was defined in dB SNR units:

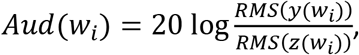

where y(*w_i_*) is the acoustic waveform of a word *w_i_* spoken by one speaker, and z(*w_i_*) is the acoustic waveform of the other speaker at the same time. Because neural responses have limited dynamic range while the audibility measure ranged from –inf to inf, the audibility values were rescaled to range from 0 to 1. In order to do this, audibility values were first clipped above 10 dB and below −10 dB, and then scaled to the 0-1 range by:

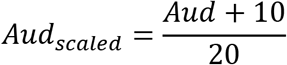

Finally, because the distributions of regressor values had distinct means for different features, we normalized each feature’s non-zero regressor values to have an RMS of 1. Bringing different features into similar amplitude ranges was done in order to make the amplitudes of corresponding TRFs more similar to each other, thus improving regularization performance.

It is notable that although neither dissimilarity, nor surprisal correlated with audibility (r = 0.03 and −0.02, respectively), there was a modest correlation between dissimilarity and surprisal (r = 0.22), suggesting that both features captured some aspects of speech predictability. Nevertheless, the fact that the correlation was relatively low suggests that much of the variance in each of the two features captured distinct aspects of the linguistic content in the speech stimuli.

#### 2.9.2 Feature-specific model performance

After fitting the full three-feature model as described above, we computed the unique contribution of each feature to the overall model fit using procedures described in Broderick et al. (2020). Briefly, on each cross-validation fold, we estimated each feature’s contribution to the overall fit by comparing the goodness-of-fit for the full model to a null model, in which that feature’s contribution was eliminated. This was done by permuting regressor values of that feature, while maintaining their original timing. For all other features, the original regressors were used. Null model fits were computed by convolving the estimated TRFs with these regressors and correlating the predicted EEG waveform with the test data. This procedure was repeated 10 times to estimate the average null-model performance. Each feature’s model contribution was then computed as the difference between the goodness-of-fit metrics for the full model and its null model.

#### 2.9.3 Regions of interest

To strengthen our statistical analyses in light of inter-subject variability due to nuisance variables such as head shape and electrode cap placement, all analyses were performed on two regions of interest (ROI) derived by averaging model goodness-of-fit and TRFs from subsets of frontal and parietal electrodes (Fig. 1B). The parietal ROI was chosen because of prior evidence that responses to higher-level features such as dissimilarity or surprisal tend to peak over parietal sites near electrode Pz (e.g., Broderick et al., 2018; Weissbart et al., 2019). The frontal ROI was included because we hypothesized that recruitment of frontal regions may aid prediction and disambiguation of the speech signals, particularly in challenging listening scenarios such as in the presence of a competing speaker.

#### 2.9.4 Statistical analysis

Group-level statistical analyses were applied to pooled outputs of single subject TRF analyses. Prior to performing statistical tests, outliers were detected using a two-stage approach, applied separately to samples from each age group to minimize the influence of true between-group differences on this procedure. First, full model goodness-of-fit values that were more than 1.5 inter-quartile ranges (IQR) below the goodness-of-fit corresponding to the lower quartile, or 1.5 IQR above the value corresponding to the upper quartile were detected as outliers. No participant met this criterion. Second, for each feature’s TRF for the attended stories (which were generally more robust compared to the ignored stories), we used the same 1.5 IQR criterion to detect outliers at each time point of the TRF. Subsequently, we computed the proportion of outlier time points for each subject. We set the outlier-proportion criterion to 0.15, so that participants with more than 15% of outlier time points were detected as outliers. This led to the exclusion of 2 participants (1 YA, and 1 OA), leaving a total of 39 participants (19 YA and 20 OA, including 17 with HL) in the analysis.

A mixed-design ANOVA with a between-subjects factor of age group (YA vs. OA), and within-subject factors of ROI (frontal vs. posterior), model feature (dissimilarity, surprisal, and audibility), and attention (attended vs. ignored story) was used to assess how these factors related to the feature-specific contributions to the model fit. Post-hoc tests were conducted using two-tailed t-tests or the analogous non-parametric test, depending on the outcome of an Anderson-Darling test of normality on the data.

Comparisons of TRFs for the attended and ignored stories were performed for each time point of the TRFs using two-tailed, paired-samples t-tests. Because this involved hundreds of statistical comparisons, we applied the *false discovery rate* (FDR; Benjamini and Hochberg, 1995) correction to control for the proportion of false positives among all significant discoveries. Similarly, between-group comparisons (i.e., younger vs. older adults) were performed on TRF time courses, with two-sample t-tests applied separately to the attended and ignored TRFs and corrected using the FDR method.

Finally, exploratory correlation analyses were performed on different combinations of neural (e.g., full model goodness-of-fit, feature-wise model contributions, TRF amplitudes) and behavioral metrics (e.g., comprehension, confidence, and SSQ_m_ scores). In these analyses we corrected each set of correlations using the Bonferroni correction. Importantly, we used less stringent multiple comparisons correction (i.e., not correcting by the total number of comparisons across all combinations of correlated variables), because of the large number of comparisons performed.

## 3. Results

### 3.1 Behavioral measures of speech understanding

Following each 1-minute block of listening to a two-talker speech mixture, participants responded to four true/false questions about the content of the attended story and indicated their confidence about their response. The average performance on this comprehension task was 83.2% (SD: 6.8%, 65.9 - 94.2% range), significantly above the 50% chance level [t(38) = 30.48, p < 0.001], indicating that participants were successfully able to attend to the target speaker and comprehend the content of the story. We found a significant effect of age on performance [t(37) = −3.04, p = 0.004], with older participants performing better than younger participants (YA: mean ± s.d. = 80.1 ± 7.5%, OA: 86.1% ± 4.6%). A correlation analysis with age used as a continuous variable showed the same association with the proportion of correct responses (r = 0.33, p = 0.043). Confidence measures showed the same general pattern of results as the comprehension scores and the two measures were positively correlated [r = 0.69, p < 0.001], indicating that participants had good awareness of their performance.

Because hearing loss was more common among the older participants, and we compensated for it by amplifying the audio in frequency ranges of elevated thresholds (see Methods), we assessed whether this amplification could account for the difference in performance. As expected, in the portion of participants who received amplification (n = 17), there was no relationship between average high-frequency audiogram (2-8 kHz range), and comprehension-performance (r = 0.06, p = 0.81) or confidence (r = 0.2, p = 0.44) measures. The same pattern was observed when using the average of the entire 0.25-8 kHz range of audiometry. As such, there was no evidence that amplification had an impact on performance, or that it could account for between-group differences in performance.

Prior to the experimental session, each participant filled out a modified subset of the SSQ (SSQ_m_) questionnaire to assess their subjective difficulties with speech-in-noise perception. We found no difference in these measures between younger and older participants (z = −0.42, p = 0.67, Mann-Whitney U-test), and no correlation between SSQ_m_ score and the proportion of correct responses from the behavioral task (r = −0.17, p = 0.29), or between SSQ_m_ and high-frequency hearing loss (r = 0.03, p = 0.91).

### 3.2 Cortical measures of speech-mixture processing

In order to characterize cortical responses to semantic content of speech, we applied computational models to EEG responses measured while participants listened to a mixture of two distinct narrative stories, while attending to one of them. The features included in the model were word audibility reflecting word-by-word fidelity of the incoming acoustic signal, semantic dissimilarity reflecting short-term (sentence timescale) dissimilarities between the word2vec vector characterizing each word and its immediately preceding context, and word surprisal reflecting long-term predictability of each word given the preceding multi-sentence context.

Linear regression of these features against the EEG signal produced responses that explained a significant amount of variance in the data pooled across participant groups and electrodes, as reflected by a significant positive correlation between the full-model EEG prediction and held-out data for both attended [t(38) = 20.87, p < 0.001] and ignored [t(38) = 8.75, p < 0.001] speech, with a significantly stronger fit for the former (t(38) = 10.60, p < 0.001; Fig. 2). The same pattern of results was observed when examining model fits in frontal and parietal ROIs. Figure 3 depicts the average attended (green) and ignored (purple) TRFs in the two ROIs for each of the features included in the model. We observed robust responses to the attended story for each of the features included in the model, with prominent early (~ 100 ms) and late (~ 400 ms) peaks in neural activity. In contrast, the ignored story elicited comparatively flatter responses, with predominantly early peaks in neural activity. Indeed, most features showed extensive periods in the early and late portions of the TRFs where attended and ignored responses differed significantly, as depicted by black horizontal bars at the bottom of each TRF plot (indicating FDR-corrected significant time points).

**Figure 2.**
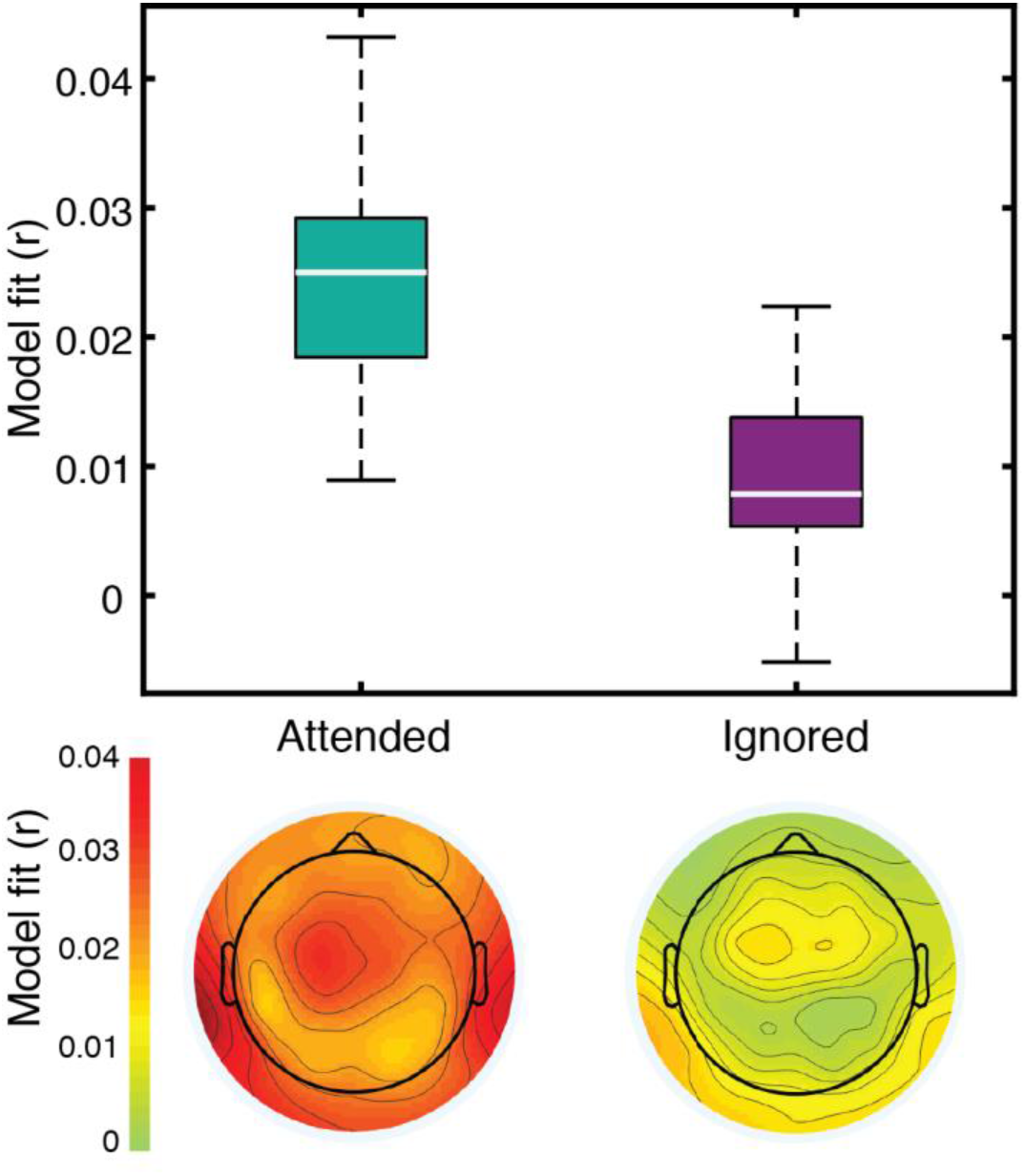
The three-feature model explained a significant amount of variance in responses to both attended and ignored speech. Box plots (top) represent distributions of goodness-of-fit values averaged over electrodes across all participants. The topographic plots (bottom) depict the distribution of goodness-of-fit values for attended and ignored speech across the scalp.

**Figure 3.**
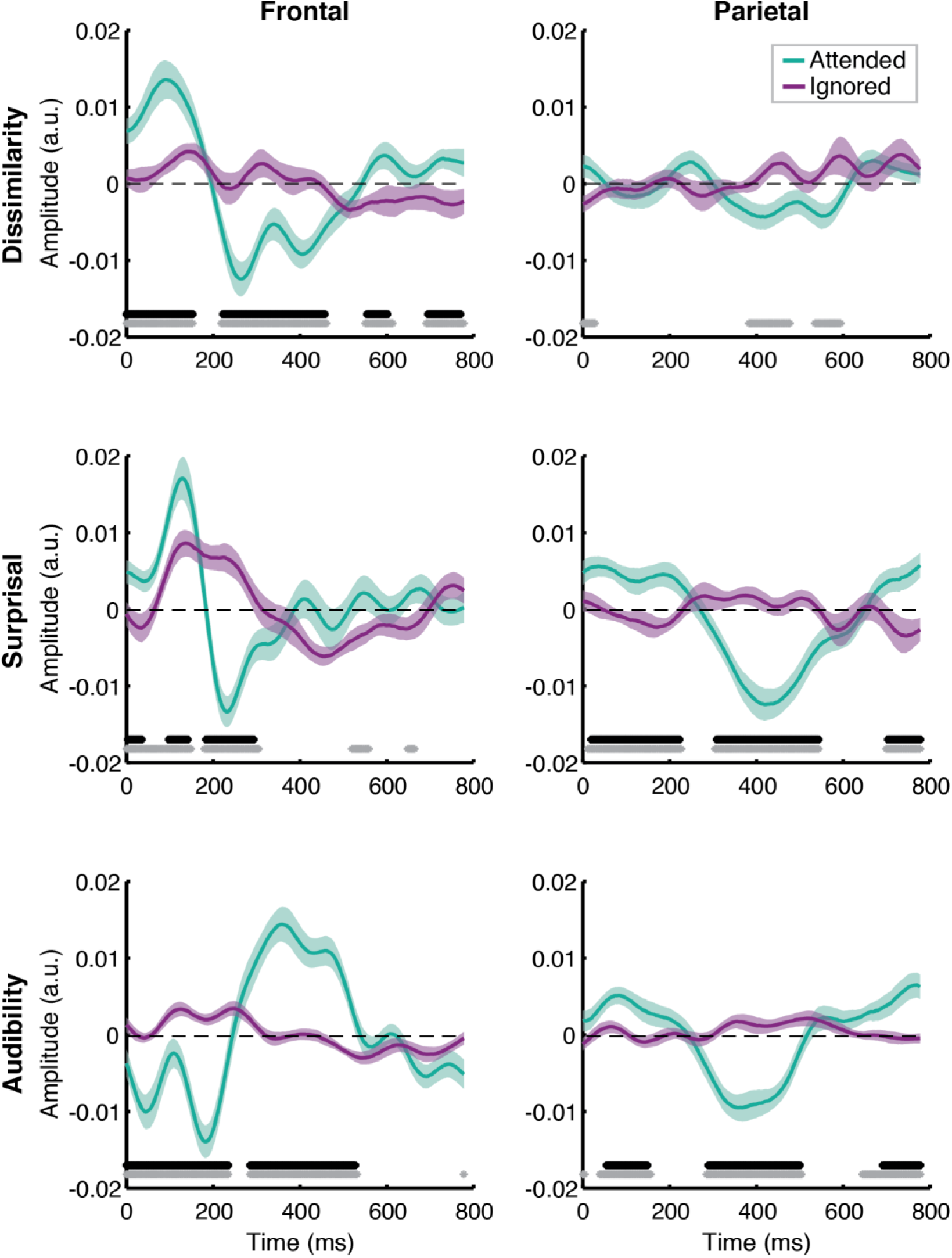
Attentional modulation of feature-specific responses. Each plot depicts the comparison of TRFs averaged across all participants for attended (green) and ignored (purple) speech for each of the features (panel rows) and ROIs (panel columns). The upper and lower bound of each curve represents ± 1 standard error (SE) of the mean. Black and gray horizontal bars at the bottom of the plots indicate time intervals over which attended and ignored TRFs differed significantly at the FDR-corrected and uncorrected level, respectively, with α = 0.05.

Contributions of each feature to the overall model fit for both age groups are plotted in Fig. 4. Model fit contribution values represent the difference in goodness-of-fit for the held-out EEG data between the full model and null models in which a given feature’s regressor was selectively disrupted by shuffling its feature amplitudes (see section 2.9.2). Thus, for a particular feature, a model fit contribution exceeding 0 represents the scenario where the EEG responses scaled, to some degree, with that feature’s regressor values. To compare how these model contributions differed in the two age groups, we performed a mixed-design ANOVA with within-subject factors of ROI, model feature, and attention, and a between-subjects factor of age group (Table 1). As expected, we found a main effect of attention [F(1,37) = 34.28, p < 0.001, *η_p_^2^* = 0.48] reflecting generally stronger tracking of high-level features within the attended than ignored speech stream. We also found main effects of ROI [F(1,37 = 8.89, p = 0.005, *η_p_^2^* = 0.19], feature [F(2,74) = 18.48, p < 0.001, *η_p_^2^* = 0.33], and age group [F(1,37 = 7.92, p = 0.008, *η_p_^2^* = 0.18].

**Figure 4.**
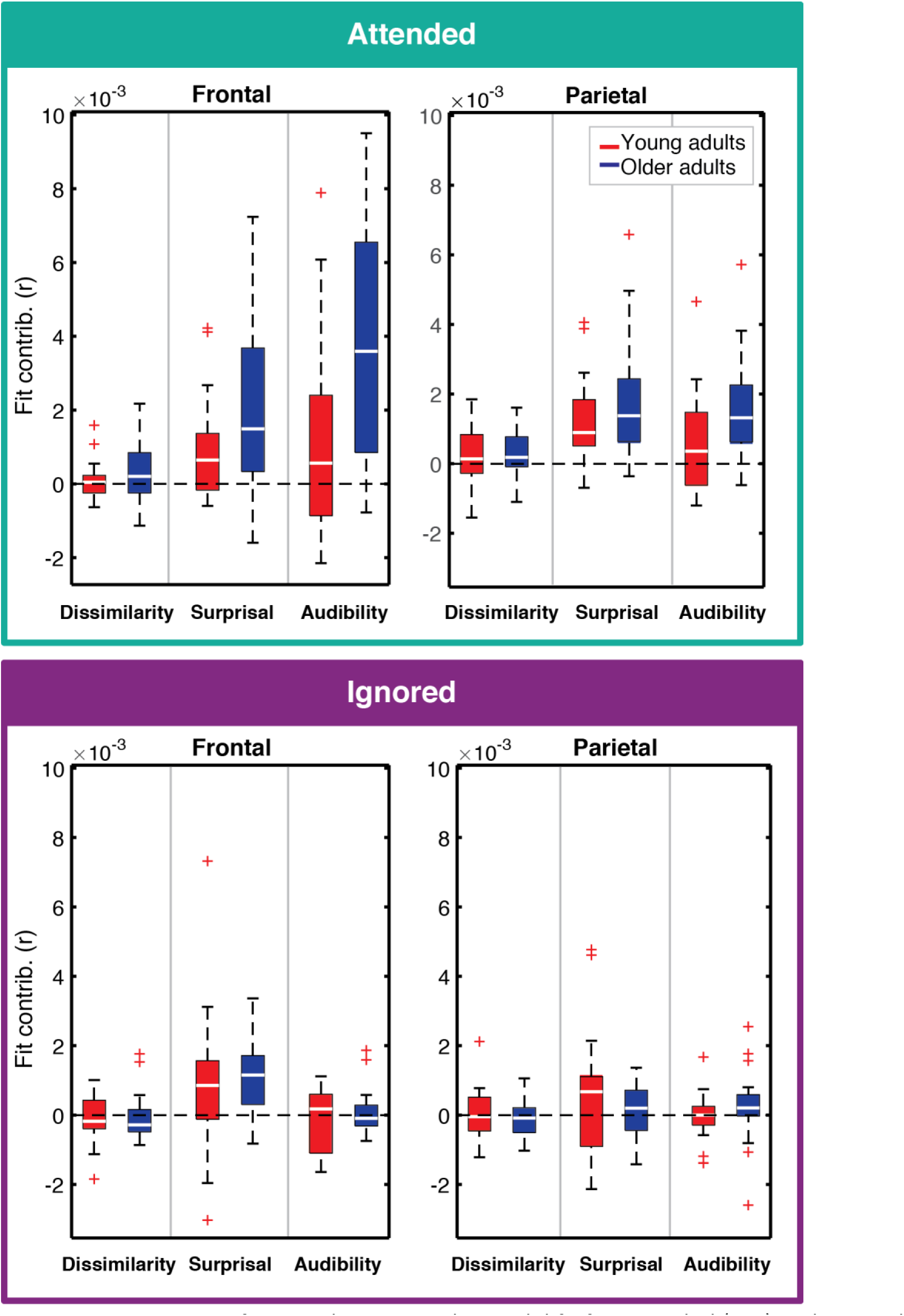
Feature-specific contributions to the model fit for attended (top) and ignored (bottom) responses. Each panel depicts the box plot of model fit contributions for each of the three features in the younger (red) and older (blue) adult groups. Left and right panels represent results for frontal and parietal ROIs, respectively. Note that some points are depicted with red + signs as outliers in order to better depict where the bulk of the points lie within the fit contribution distributions. However, all data points were utilized in statistical analyses described in the text.

**Table 1.**
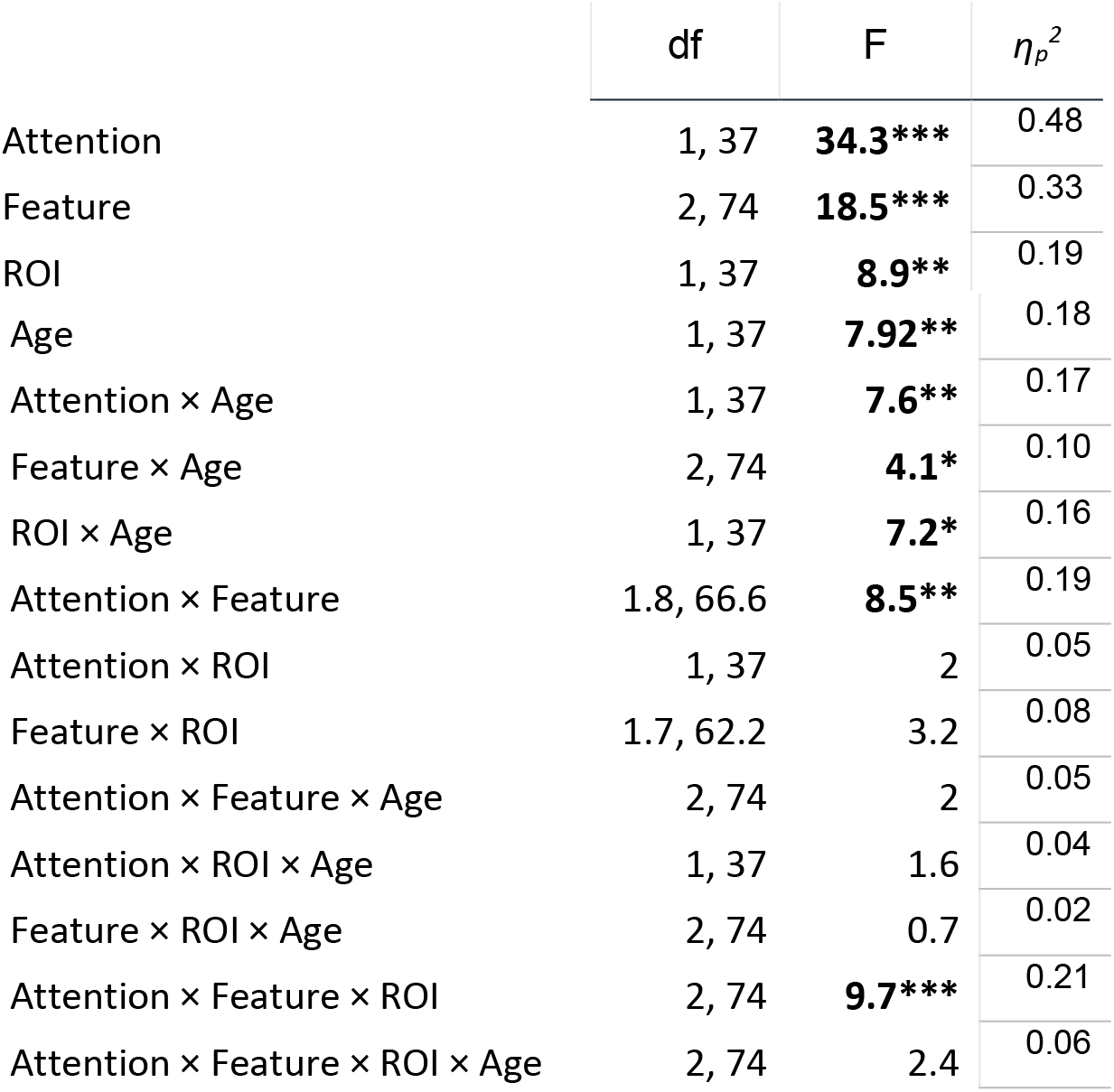
Mixed-factors ANOVA results. Significant F-statistic values are bold, with levels of significance: * p < 0.05, ** p < 0.01, *** p < 0.001

In addition to these main effects, we detected a number of significant interactions. There was a significant interaction between attention and age group [F(1,37 = 7.64, p = 0.009, *η_p_^2^* = 0.17], reflecting an overall greater difference between attended and ignored fits in older than younger participants [t(37) = −2.76, p = 0.009]. A significant interaction between ROI and age group [F(1,37 = 7.24, p = 0.011, *η_p_^2^* = 0.164] was associated with significantly stronger contributions to model fits across features at the frontal compared to the parietal ROI in older adults (p = 0.007; Mann-Whitney U-test). Third, we found a significant interaction between feature and age group [F(2,74 = 4.09, p = 0.021, *η_p_^2^* = 0.10], and a post hoc analysis revealed this was due to greater difference in contributions to model fit between word audibility and dissimilarity in older than younger participants [t(37) = −3.01, p < 0.005; Bonferroni corrected with α = 0.017].

Several interactions did not involve age group, including a significant interaction between attention and feature [F(1.8,66.63) = 8.55, p = 0.001, *η_p_^2^* = 0.19], a trend towards an interaction between feature and ROI [F(1.68, 62.18] = 3.2, p = 0.056, *η_p_^2^* = 0.08], and a three-way interaction between attention, feature, and ROI, [F(2,74 = 13.05, p < 0.001, *η_p_^2^* = 0.21]. Because the latter interaction was a combination of factors from the former two, we only pursued post hoc analyses for the three-way interaction. These indicated that in the frontal ROI, the contribution of audibility to the model fit was greater for the attended than the ignored story, and that this differential was greater than that for both dissimilarity and surprisal [t(38) = −3.38, p = 0.002, and t(38) = −3.61, p < 0.001, respectively; Bonferroni corrected with α = 0.017]. Comparison of dissimilarity and surprisal showed no difference [t(38) = 1.38, p = 0.18].

Although the goodness-of-fit analyses above indicate that there are significant differences in processing of attended and ignored speech between younger and older participants, they do not provide insight into the timing and amplitude of the underlying neural responses. To explore if our data contain evidence of age-related differences in neural responses, we statistically compared TRF amplitudes between the two age groups at each time point in the 0 to 800 ms range. Because these analyses involved hundreds of point-by-point comparisons between groups, we corrected for false discovery rate (FDF), and focused on comparisons at the level of individual features, rather than utilizing more complex interaction metrics. As such, these analyses were relatively rudimentary, and should be considered as exploratory in their nature.

Figure 5 depicts the differences in responses to the attended speech between younger (red lines) and older (blue lines) participants, separately for each feature (plot rows) and ROI (plot columns). Two-tailed statistical outcomes at the p < 0.05 level are depicted at the bottom of each plot in both uncorrected (gray horizontal bars) and FDR-corrected (black horizontal bars) forms. At the FDR-corrected level, we only found two clusters of significant time points in the frontal TRFs for dissimilarity, with older participants showing a significantly more negative response between approximately 260-300 ms, and a significantly more positive response in the 620-675 ms time range. While surprisal and audibility showed no robust differences at the FDR-corrected level, several clusters of time points were suggestive of group differences at the level of uncorrected statistics. For surprisal, we found that older adults had a greater negative deflection in the 225-260 ms time range and a pair of positive deflections around 390-430 and 515-580 ms that were absent in the TRF of the young adults at the frontal ROI. We also found a single cluster of time points with greater negative deflection for older than younger adults between 395-435 ms in the parietal ROI. For word audibility, we found a prolonged elevated response with portions between 415-480 ms exhibiting larger positive deflection in older than younger participants, at the frontal ROI. Older adults also showed a greater negative deflection in the word-audibility TRF frontally, and a greater positive deflection parietally around 550-600 ms.

**Figure 5.**
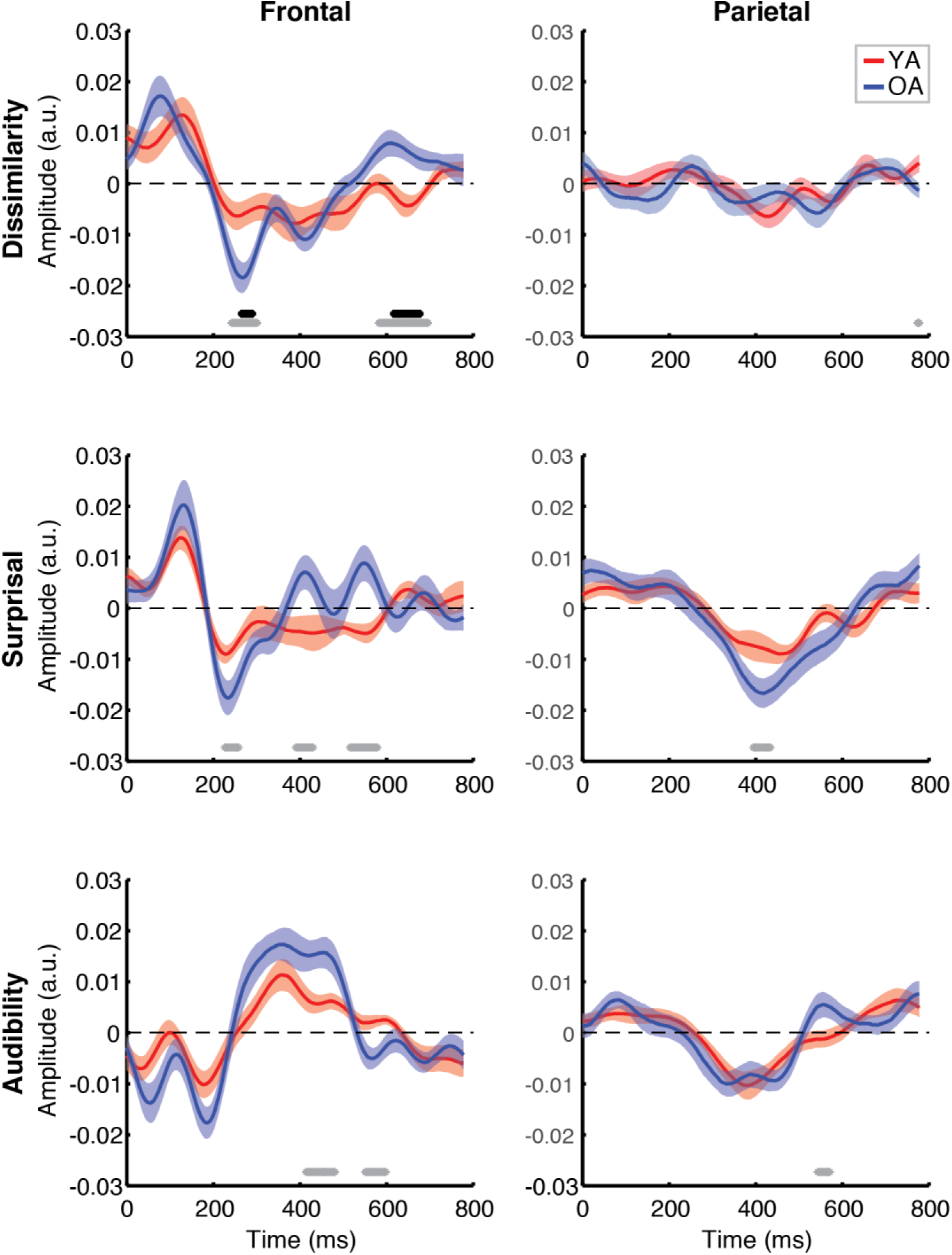
Between-group comparison of TRFs for attended speech. Each plot depicts a comparison of TRFs between younger (red curves) and older (blue curves) participants, for different features (panel rows) and ROIs (panel columns). Black and gray horizontal bars at the bottom of the plots indicate time points at which the two age groups differed significantly at the FDR-corrected and uncorrected level, respectively, with α = 0.05.

Between-group comparison of TRFs for ignored speech are shown in Figure 6. Unlike responses to attended speech, most features, with the exception of frontal TRFs for surprisal, show largely flat response patterns that do not differ between groups. Several time points showed a difference in uncorrected statistics for each of the features, the most notable of which was a more negative response of younger adults to audibility between 590-660 ms in the frontal ROI. However, given the low amplitude of the TRFs, and long latencies of most of the potential differences, we believe these are likely to simply reflect false discoveries due to hundreds of comparisons. Indeed, fewer than 5% of comparisons for ignored speech were significant at the uncorrected level.

**Figure 6.**
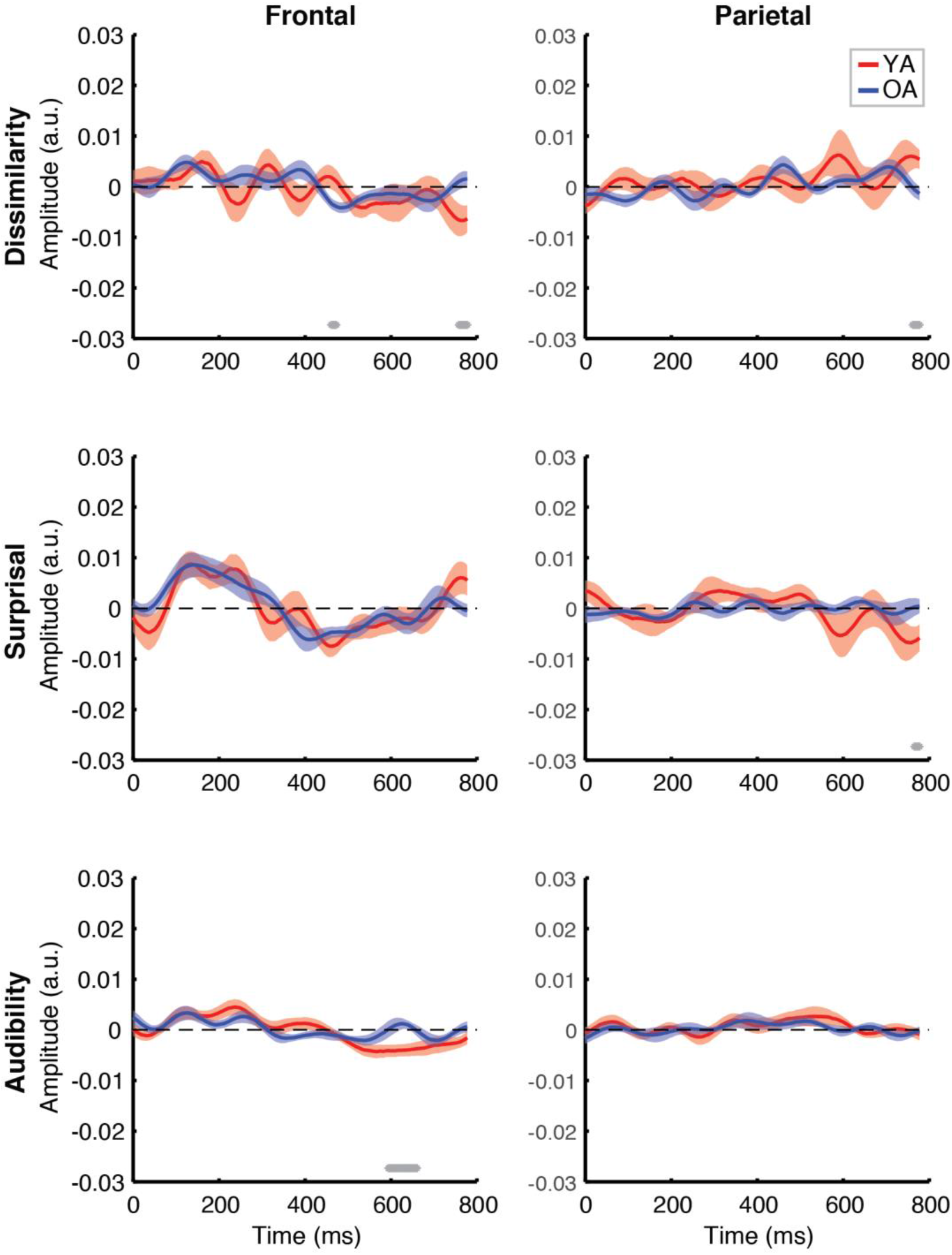
Between-group comparison of TRFs for ignored speech. Subplot arrangement and statistical comparisons are as in Fig. 5.

To complement these exploratory point-by-point analyses, we also conducted between-groups analyses specifically targeted at comparing responses in the time range of the N400 response. To this end, we compared each feature’s average TRF amplitudes in the 300-500 ms range. Because previous work found little to no evidence of N400 for ignored speech, these comparisons were only done for attended speech. Although we found both a significantly more negative parietal N400 for the older group to surprisal [t(37) = 2.03, p = 0.05], and a significantly elevated frontal response in the older group for audibility [t(37) = −2.72, p = 0.01], neither of these results remained significant with Bonferroni correction (α = 0.008, given the total number of 6 comparisons).

### 3.3 Neuro-behavioral correlations

We next sought to examine how our electrophysiological measures related to behavioral responses during the experiment, and the SSQ_m_ scores obtained prior to this experiment. To this end, we conducted a number of exploratory analyses, including correlations between behavioral measures and the overall model goodness-of-fit, feature-specific model contributions, and the average TRF amplitudes in the 300-500 ms time range. Given the number of these analyses, and our limited sample size, we focused our analyses on full participant samples, rather than age group comparisons. Because of the less stringent multiple comparisons correction procedure (only correcting by the number of statistical tests within each analysis), significant effects in this section should be interpreted as trends rather than true statistical effects.

Figure 7 depicts the relationship between the proportion of correct responses on comprehension questions during the experiment, and the overall model goodness-of-fit in the frontal (left panel) and parietal (right panel) ROIs. While we observed no relationship in frontal regions (r = 0.21, p = 0.19), there was a marginally significant positive association between the two measures (r = 0.35, p = 0.027, Bonferroni corrected α = 0.025) in the parietal ROI. A similar pattern of results was observed when average confidence ratings for the comprehension questions were used instead of the performance itself. Relationships between the proportion of correct responses and feature-specific contributions to the model fit are depicted in Figure 8. We observed a trend towards a positive association for word audibility in both the frontal (r = 0.29, p = 0.07) and parietal ROIs (r = 0.4, p = 0.011), although neither correlation reached significance after correcting for multiple comparisons (α = 0.008). None of the other features showed a significant association with comprehension scores.

**Figure 7.**
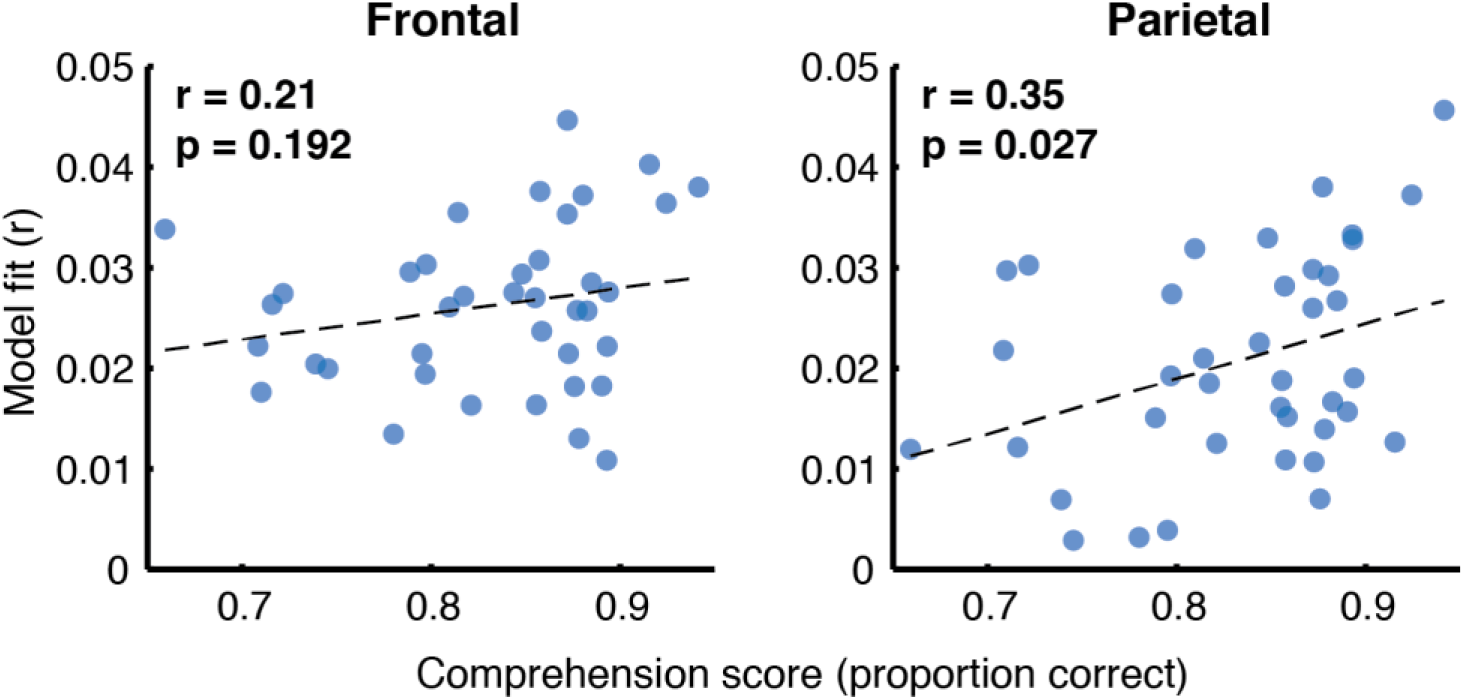
Scatterplots showing the relationship between the full model goodness-of-fit and the proportion of correct responses on the comprehension questions. Pearson’s correlation coefficients and the corresponding uncorrected p-values are shown for frontal (left plot) and parietal (right plot) ROIs. Symbols represent data from individual participants pooled across the two age groups, YA and OA.

**Figure 8.**
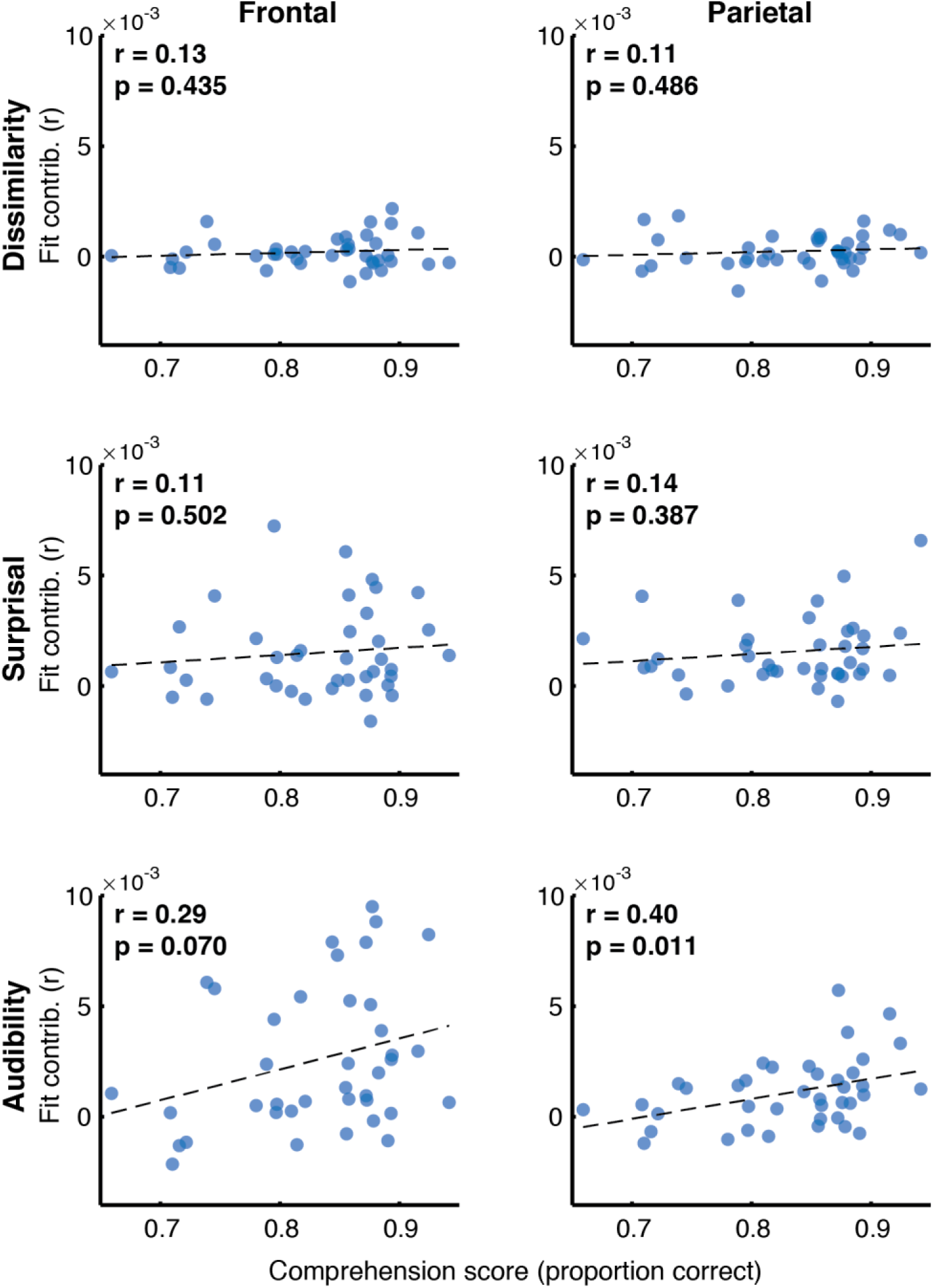
Scatterplots of comprehension scores and feature-specific model contributions. Different rows of panels refer to different features and different columns correspond to the two ROIs. Pearson’s correlations and the corresponding uncorrected p-values are shown in the upper portion of each panel.

Next, we explored the possible relationship between the comprehension scores (proportion correct) and the average TRF amplitude in the 300-500 ms time range, when N400 effects generally appear parietally. These analyses, shown in Figure 9, revealed trends towards a positive relationship in frontal regions for surprisal (r = 0.33, p = 0.037) and audibility (r = 0.31, p = 0.059), as well as a trend towards a negative relationship for surprisal in parietal ROI (r = - 0.29, p = 0.075). As before, none of these associations were significant when correcting for multiple comparisons. Although this analysis focused broadly on the time range of N400, two of the frontal trends were associated with positive, rather than negative deflections in the TRF.

**Figure 9.**
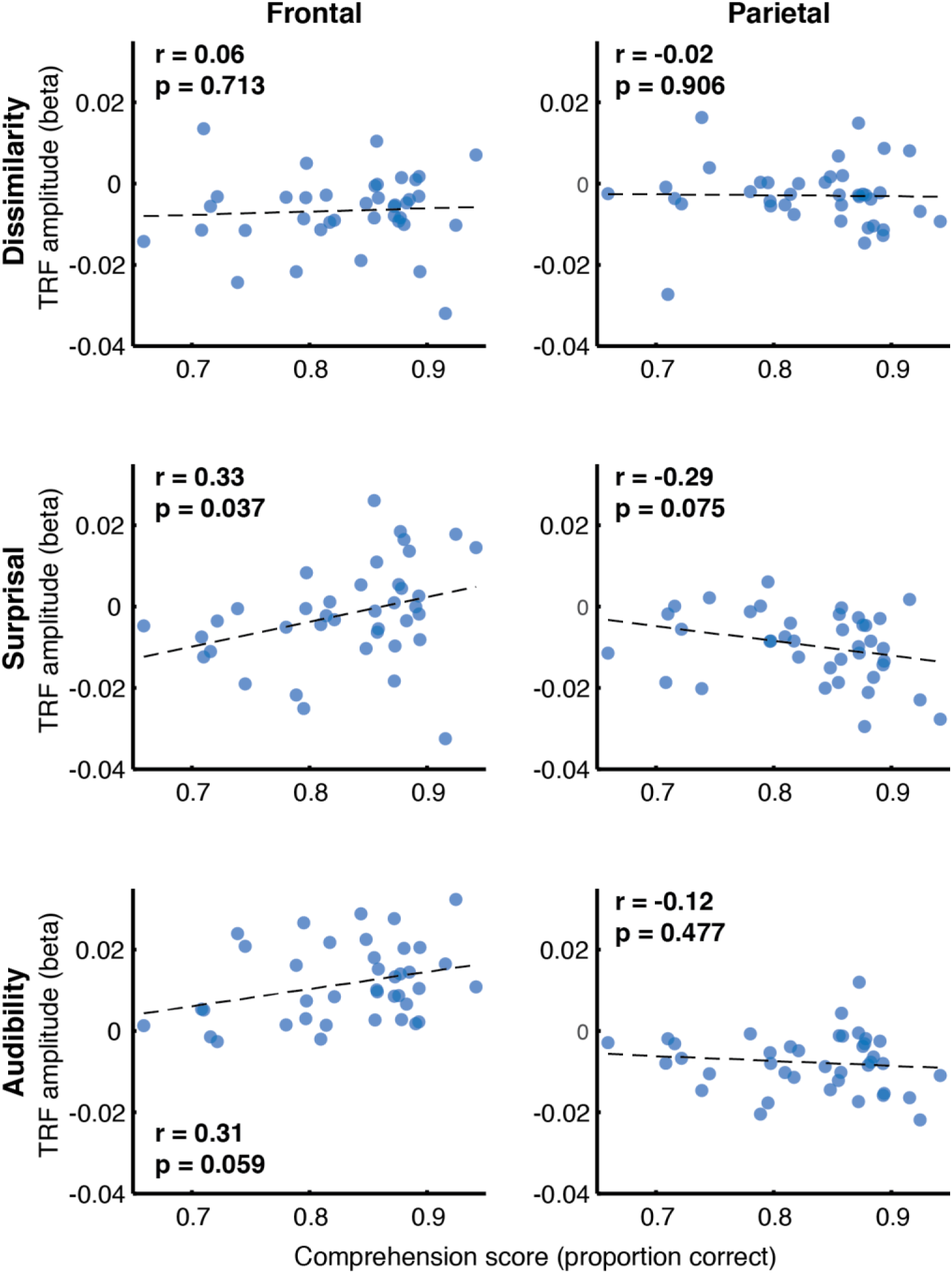
Scatterplots of comprehension scores and mean TRF amplitudes between 300-500 ms. Figure layout is as in Fig. 8.

Correlation analyses examining the relationship between subjective SIN perception difficulties, captured by the SSQ_m_ scores, and the full model goodness-of-fit metric (Fig. 10) revealed trends towards a negative relationship in both the frontal (r = −0.30, p = 0.064) and parietal ROIs (r = −0.32, p = 0.044). However, analyses of relationships with feature-specific TRF amplitudes and model contributions revealed no feature for which these trends were apparent.

**Figure 10.**
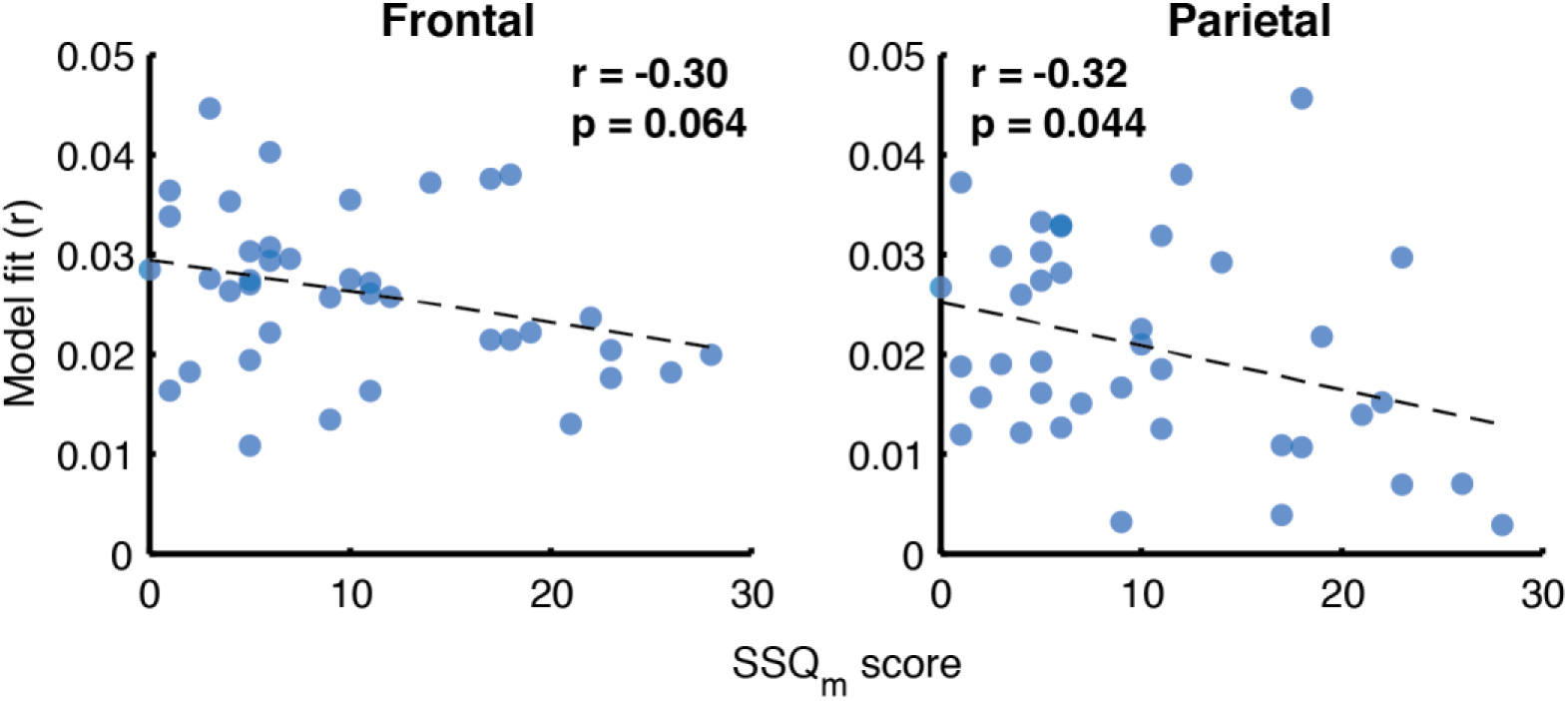
Scatterplots of SSQ_m_ scores and overall model goodness-of-fit for frontal (left panel) and parietal (right panel) ROIs. Note that a higher score on SSQ_m_ questionnaire reflects a greater difficulty with understanding speech in noise.

Finally, because a portion of the participants had mild hearing loss at high frequencies (which was compensated for by amplifying speech in the corresponding frequency ranges; see Methods), we examined if and how high-frequency (2-8 kHz) hearing thresholds related to the overall model fits (Fig. 11). Although we found no relationship between the average hearing thresholds over the 2-8 kHz range and model goodness-of-fit for attended speech (Frontal ROI: r = −0.04, p = 0.87; Parietal ROI: r = −0.02, p = 0.95), there was a significant negative correlation for ignored speech both frontally (r = −0.59, p = 0.013) and parietally (r = −0.62, p = 0.008). At the level of feature-specific contributions to the model fit, there was no indication that this negative correlation was driven by any particular feature, as most features showed low, non-significant negative correlations.

**Figure 11.**
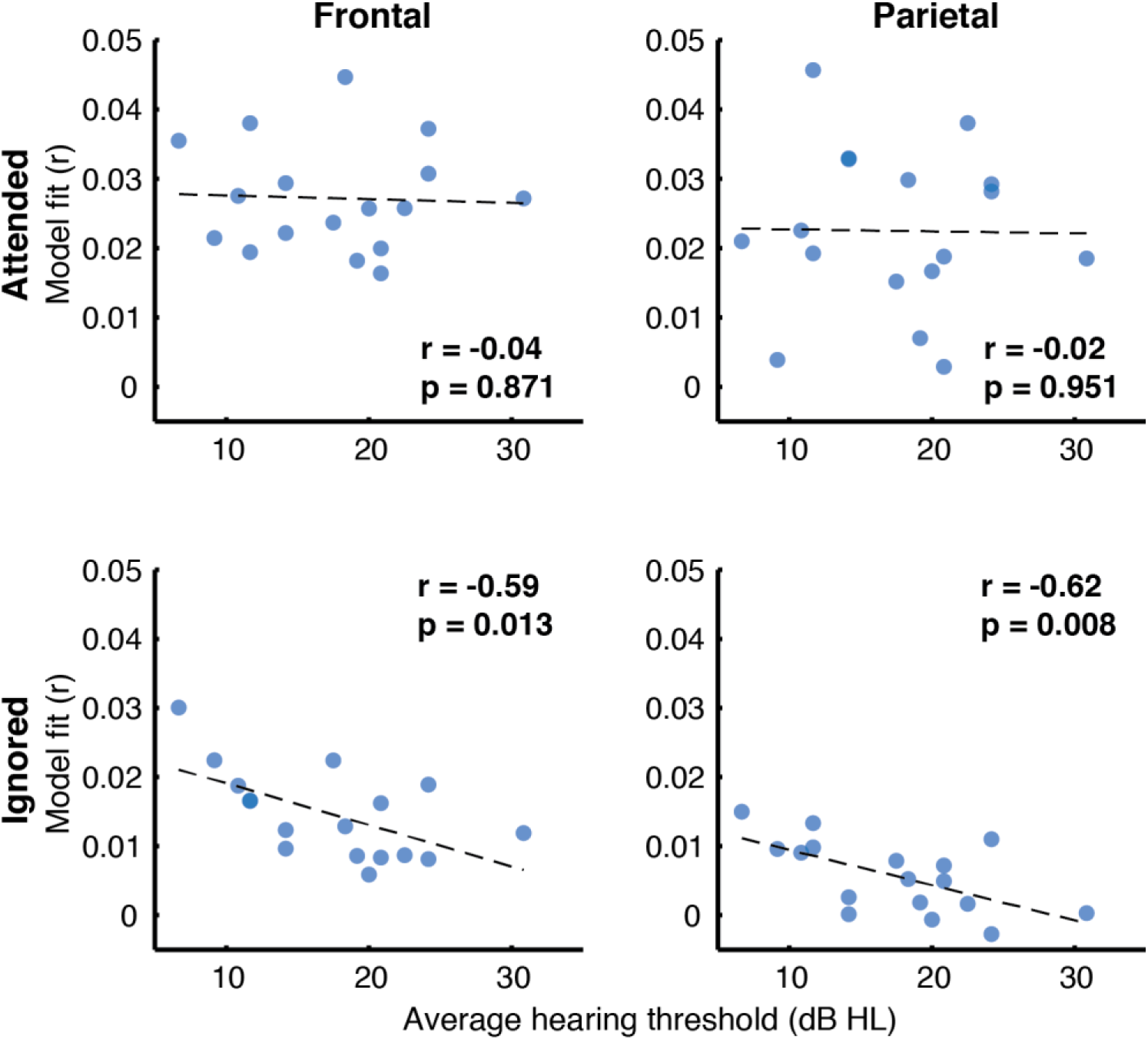
Scatterplots of average high-frequency hearing thresholds (2-8 kHz) and overall model goodness-of-fit as a function of attention (panel rows) and ROI (panel columns).

## 4. Discussion

Speech perception is a fundamental capability of the human auditory and language systems, facilitating our abilities to learn and engage in various types of social interaction. However, deficits in SIN perception are commonly experienced by the aging population (e.g., van Rooij and Plomp, 1990; Goossens et al., 2017) and are reported surprisingly frequently even among the younger and nominally normal hearing population (Saunders, 1989; Zhao and Stephens, 2007; Tremblay et al., 2015). Importantly, while subjective SIN perception difficulties may indicate a significant adverse impact on quality of life (Dalton et al., 2003; Chia et al., 2007), existing objective (laboratory and clinical) measures of speech perception have shown surprisingly poor correlations with the self-reported difficulties as measured, for example, by SSQ scores (Phatak et al., 2018; Smith et al., 2019).

In the present study, we measured EEG responses to continuous two-talker speech mixtures in younger (< 40 y.o.) and older (> 40 y.o.) participants. Participants’ cortical responses in the 1-8 Hz range were predicted by modeling TRFs for three speech features, short-timescale semantic dissimilarity, long-timescale lexical surprisal, and word-level audibility. We also collected behavioral measures, including participants’ subjective ratings of their difficulties with SIN understanding (modified SSQ), and comprehension scores for attended speech during the experiment and the associated confidence ratings.

Our three-feature model was able to explain significant variance in the EEG data, especially in responses to attended speech, where each of the features contributed to the neural responses (Fig. 4). The evidence for this was particularly strong for surprisal and audibility, suggesting that these model features captured stimulus characteristics that were actively tracked by our participants’ auditory systems. Moreover, we found that participants’ performance on the comprehension task (Fig. 7), as well as the associated confidence ratings, showed a trend towards a positive correlation with the goodness of the overall model fit for the attended speech, suggesting that successfully tracking these features is related to speech comprehension. Although our data does not support a strong association between performance and model contributions, or TRF magnitudes, for any one of the model features, we did find trends towards an association between word audibility and performance in both ROIs (for both model fit contributions, and TRF magnitudes), and in the frontal region between the surprisal TRF magnitude and performance. Speculatively, these trends suggest that improved comprehension may be related to at least two cognitive processes. First, the association with audibility suggests that improved performance may stem from more effective weighing of word-level information by word reliability, as reflected by the word SNR. Second, the association with surprisal suggests that high performance may be related to increased sensitivity to lexical and/or semantic associations between different segments of speech.

Consistent with previous work on neural representations of two-talker speech (Ding and Simon, 2012; Mesgarani and Chang, 2012; Broderick et al., 2018; O’Sullivan et al., 2019) we found robust differences between responses to attended and ignored speech both in the goodness of model fits and the TRFs. In general, model fits were better for attended than ignored speech (Fig. 4) and the associated TRFs for attended speech showed complex, multi-peaked morphologies, whereas the responses to ignored speech were flatter and contained fewer prominent peaks (Fig. 3). Thus, our results indicate that responses to a speech mixture preferentially reflect attended speech, while representations of distractor speech are largely suppressed.

Comparisons of EEG responses between age groups revealed a complex pattern of age-related differences, captured particularly by model fit measures. Specifically, we found that older participants exhibited on average greater differences in feature-specific model-fit contributions between attended and ignored speech. This age effect was driven primarily by better fits for attended speech in the frontal ROI (see Fig. 4). Although to a weaker degree, these differences were mirrored in attended TRFs, in that older adults showed generally stronger TRF deflections from 0 compared to younger participants (Fig. 5). In most cases, however, these TRF differences did not reach statistical significance when controlling for false discovery rate, possibly due to nuisance factors such as inter-subject variability in cortical geometry, and/or inadequate sample size.

With respect to the modelled features, we found that surprisal and audibility both showed stronger frontal contributions in older adults, whereas parietal contributions were relatively similar between the two groups. We speculate that the stronger fits in the frontal region in older adults may be indicative of heightened reliance in this group on both lexical prediction, as reflected by increased accuracy of surprisal fits, and on words with better audibility. Higher word SNRs may have been more important for disambiguation of the masked portions of speech for older compared to younger adults. Although audibility itself reflects a relatively low-level aspect of our stimuli, its frontal TRF profile showed a prolonged positive deflection in the 250-550 ms latency range. Such a long latency is consistent with the possibility that this audibility-related response may reflect engagement of higher-level processes, such as retrospective disambiguation, or prospective prediction.

It is notable that participants in the older group exhibited significantly better performance on the comprehension task than younger adults, despite having greater prevalence of hearing loss (15 out of 17 participants with HL were in the older group and the degree of HL was not significantly correlated with performance). This difference in performance difference complicates the interpretation of age-related differences in neural responses. It may be the case that older adults in our participant sample were either more engaged, or exerted greater effort in the task, which in turn led to stronger speech tracking in their EEG data, as well as better performance. This is plausible, since more participants in the older group (12/20 older vs 8/19 younger participants) indicated having a subjective sense of experiencing greater difficulty with SIN understanding compared to their peers. The sense of greater difficulty may have motivated at least some of the older participants to exert greater effort to perform well. However, while the average performance of participants with self-reported SIN difficulties was slightly better than that in participants who did not report such difficulties, these differences were not significant. Despite this, the possibility that differences between the two age groups in effort, attentiveness, or another factor may underlie the neural differences discussed above, deserves further attention in future work.

### 4.1 Relationship to existing work on age-effects on electrophysiological measures of speech processing

Several studies have examined effects of age (Presacco et al., 2016; Decruy et al., 2019; Zan et al., 2020) and hearing loss (Millman et al., 2017; Decruy et al., 2020) on continuous speech processing in the context of envelope tracking. Generally, these studies have demonstrated that older adults and those with hearing loss exhibit exaggerated cortical tracking of speech envelope both in quiet and in the presence of a competing speaker, as reflected by higher envelope reconstruction accuracies from delta-band EEG or MEG responses in these populations. Our analyses show a similar pattern of amplified feature tracking in the aging population, albeit for word-level features. Responses to the audibility feature, in particular, may reflect similar underlying processes as those involved in envelope processing. However, audibility in our study was defined as the word-by-word ratio between the acoustic energy in the two speech waveforms, rather than the absolute amplitude of each speech signal, making direct comparisons of the two measures difficult. Distinct from envelope TRFs, the audibility TRF in our study contained prolonged deflections from 0 in the 300-500 ms latency range, suggesting that our measure may tap into additional higher-level processes. Although lexical surprisal is seemingly unrelated to speech envelope, it is possible that predictive processes may interact with lower-level stimulus encoding via feedback processes, as has been demonstrated for dissimilarity (Broderick et al., 2019).

While measures of envelope tracking have provided important insights into speech processing, they are largely uninformative about the nature of higher-level processes involved in speech perception. In recent years, an increasing number of studies have investigated the relationship of electrophysiologically-measured cortical responses to both intermediate speech representations such as those evoked by different phoneme categories (Di Liberto, 2015; Lesenfants, 2020; Teoh & Lalor, 2020; but cf. Daube et al. 2019) or phonotactics (Di Liberto, 2019), and word-level representations related to lexical (e.g., Brodbeck et al., 2018), as well as syntactic and semantic (Broderick et al., 2018; Weissbart et al., 2019; Heilbron et al., 2019; Donhauser & Baillet, 2020) processing. Nevertheless, relatively little is known about how these representations change as a function of age, particularly in challenging listening conditions. Recently, Broderick et al. (2020) compared representations of semantic dissimilarity and 5-gram lexical surprisal derived from responses to clean speech in younger and older adults. They showed that although younger adults exhibited robust responses to each feature, older adults only showed strong responses to lexical surprisal (albeit with a delayed peak response), with a nearly absent response to semantic dissimilarity. These results were interpreted as potentially reflecting lesser reliance of older adults on semantic predictive process, thought to be captured by the dissimilarity feature, due to age-related cognitive decline. Consistent with this, older participants with greater semantic verbal fluency, a measure related to the ability to engage in semantic prediction, showed greater contribution of semantic dissimilarity to the model of cortical responses to speech.

Because our experimental design involved listening to a more challenging, two-speaker mixture, direct comparisons of our results with those of Broderick et al. (2020) are not possible. Nevertheless, there are marked differences between the patterns of results observed in their study compared to ours. In particular, we observed stronger tracking of both lexical surprisal and word audibility in older than younger adults, and generally weak but otherwise similar tracking of dissimilarity in the two groups. Notably, this was observed predominantly at the frontal ROI, with the posterior ROI showing a smaller difference (albeit in the same direction as the frontal results). In contrast, Broderick et al. focused their analyses on posterior electrode sites, making it unclear how tracking of their features behaved at more frontal sites that are involved in tasks relying on working memory (e.g., Gevins et al., 1997; Onton et al., 2005).

In Broderick et al. (2020), the greatest age-related differences were shown for semantic dissimilarity, whereas our goodness-of-fit results showed relatively weak contributions from this feature (compared to surprisal and word audibility) that did not differ significantly between the younger and older age groups. However, we did observe greater frontal TRF deflections in the older group for dissimilarity, with significant group differences around 250 and 600 ms, suggesting an increased gain for this feature in the older population. This underscores the importance of analyzing both model fits and the corresponding TRFs, as morphological differences in the latter may be possible even in the absence of differences in the model goodness-of-fit. The most notable difference in our results with respect to dissimilarity is that we did not observe posterior N400 response in either group, in contrast to the significant parietal N400 in the TRF for dissimilarity in older but not younger adults reported by Broderick et al. Although this discrepancy is puzzling given the use of nearly identical methods for computing dissimilarity, it raises the possibility that the utility of dissimilarity may be limited if other features, which better capture neural responses that would otherwise be attributed to dissimilarity, are included in the model.

Another important difference between the two studies pertains to the role of surprisal in the models fitted to the data. Specifically, unlike the relatively simple 5-gram surprisal used by Broderick et al., which was intended to capture responses related to the knowledge of word co-occurrence within 5-word neighborhoods, the surprisal features utilized in our study were computed using an advanced natural language model (GPT-2; Radford et al., 2019) that uses preceding context of up to several hundred words (i.e., dozens of sentences) in order to estimate each upcoming word. As such, surprisal in our study likely captured responses related to higher-level lexical and/or syntactic predictions. Thus, although responses to these two surprisal measures cannot be directly compared, the stronger tracking of surprisal by older adults in our study is consistent with increased reliance on predictive processes in this population. This is in agreement with behavioral results demonstrating greater reliance on semantic context in populations with compromised representations of speech, such as those with hearing loss (Benichov et al., 2012; Lash et al., 2013) and cochlear implants (Amichetti et al., 2018; Dingemanse and Goedegebure, 2019; O’Neill et al., 2019).

Importantly, the seemingly conflicting pattern of results between these studies could in fact reflect two distinct contributors to speech perception difficulties in older adults, namely decreases in the fidelity of lower level representations, and cognitive decline. Prevalence of mild high-frequency hearing loss in our sample of older adults was quite high, making it likely that decreased fidelity of peripheral representations had an effect on our results. While Broderick et al. did not report audiogram measures for their sample of older adults, the mean age was considerably greater in their study (mean ± s.d. = 63.9 ± 6.7 years vs 53.5 ± 8.7 years in this study), making it likely that similar or greater hearing difficulties may have impacted their participants. However, because of the age difference in the two samples, the effects of cognitive decline may have contributed more significantly to the results of Broderick et al., and may potentially explain why measures related to predictive processes showed opposite effects in the two studies. This exemplifies the complex combination of etiologies that may underlie speech perception difficulties, and the distinct ways in which they may affect speech processing. Future work should attempt to quantify these factors and use multivariate analyses to better characterize if and how they may relate to different neural measures of speech processing.

### 4.2 Higher-level speech feature tracking as an index of speech in noise perception difficulties

A key reason for our choice to study responses to lexical and semantic features is their potentially greater sensitivity to SIN perception difficulties, compared to responses driven by lower-level features such as the speech envelope. Specifically, because dissimilarity and surprisal (but not audibility) depend on preceding lexical and semantic context, in order for language processing mechanisms to accurately track them, each word within the sequence needs to be recognized and integrated with the preceding context. Lower-level SIN processing impairments may thus disproportionately impact tracking of these features. This is because missing a given word may potentially distort neural computations of surprisal and lexical predictions for a large number of subsequent words. This distortion could result in a mismatch between the objectively computed sequences of these features (used in the model) and their internal estimates.

Dissimilarity, in particular, depends on local word context (limited to one sentence, in our model). Misperception of individual words may thus greatly distort the internal estimates of the semantic relationships between words within this short-term context, leading to poor correspondence with the objectively computed dissimilarity values. Spectrally degraded speech has previously been shown to elicit weaker N400 responses, and a reduced difference in N400 between sentences with high and low cloze probabilities (Aydelott et al., 2006; Obleser and Kotz, 2011; Carey et al., 2014). Similarly, our results showed weak model contributions of dissimilarity with N400 responses essentially absent in the posterior ROI, consistent with the possibility that challenging listening scenarios may indeed disrupt representations related to relationships between words in a local context. Notably, however, we did not observe a reliable association between individual differences in the tracking of this feature, or the magnitude of N400, and performance on the comprehension task, the associated confidence measures, or the SSQ_m_. As such, the magnitude of dissimilarity tracking, or the associated TRFs, may not actually reflect the degree of SIN perception difficulties, as we hypothesized it would. Thus, it is possible that weak tracking of dissimilarity in our study may reflect that dissimilarity, as computed here, is a relatively unimportant feature for characterizing cortical speech processing. Note that although our results appear to be at odds with Broderick et al. (2018), who demonstrated robust dissimilarity-related N400 responses for both clean and two-talker speech, that study used dissimilarity as the sole feature. It is, therefore, possible that their estimated TRFs may have captured contributions from other features time-locked to word onsets (e.g., ones related to lexical and syntactic processing). Indeed, in a recent reanalysis of cocktail party data from Broderick et al. (2018), Dijkstra et al. (2020) showed that replacing dissimilarity values in a regressor with unit-amplitude impulses leads to estimation of essentially identical TRFs to those obtained with the impulses scaled by dissimilarity features. This insensitivity to impulse scaling calls into question the extent to which said TRFs reflect dissimilarity-related processing. Comparisons of single-feature TRFs derived from our data using word onset and dissimilarity regressors (analyses not shown here) mirrored these observations, suggesting that the utility of dissimilarity in explaining EEG responses to continuous speech may be limited.

In contrast to dissimilarity, our observation of robust model contributions and posterior N400 responses for surprisal suggests that this feature may be relatively robust to challenging listening scenarios. This may be the case because surprisal, as defined in the present study, reflects predictability of each word given a multi-sentence preceding context (vs. single-sentence context for dissimilarity), potentially making misperception of individual words have relatively low impact on lexical predictions. In other words, failure to recognize individual words may have a relatively small impact on the internal predictions, as these may be highly constrained in natural speech by the successfully identified words within the longer-term context. Admittedly, the apparent robustness of surprisal to adverse listening conditions may be specific to longer narratives where long-term semantic dependencies exist, such as audiobooks used in our study. In contrast to dissimilarity, we did observe weak trends suggesting an association between the amplitude of the surprisal TRF in the N400 latency range, and the performance on the comprehension questions. As such, it is possible that surprisal responses may indeed reflect the extent of SIN perception difficulties. However, because these trends were not statistically robust to multiple-comparisons correction, and because similar trends were not observed for SSQ_m_, it remains unclear if this neuro-behavioral association is reliable. A replication study with a larger sample size, improved EEG denoising algorithms, and/or more sensitive behavioral measures may be needed to further explore this link.

It is notable that the correlations between SSQ_m_ or task performance and feature-specific model contributions were overall relatively weak in this study. Although this implies that none of the features utilized in our study can on their own predict the degree of SIN perception difficulties, it is possible that such deficits may be better characterized in terms of a multi-dimensional pattern of feature-specific neural responses. In other words, it may be the case that in order to predict the extent of SIN perception difficulties, a combination of neural measures across multiple lower- and higher-level speech features needs to be taken into account. Along these lines, Lesenfants et al. (2019) showed that speech reception thresholds can be predicted from EEG responses to speech more accurately using a model that contains both spectrogram and phonetic features, compared to models containing only one of the features. Furthermore, because SIN perception difficulties can have different underlying etiologies, with different relative contributions from peripheral damage and cognitive factors, it may be the case that distinct patterns of feature-specific responses characterize different underlying causes of SIN deficits.

### 4.3 Behavioral correlates of self-reported SIN difficulties

Our data revealed a trend towards a negative association between SSQ_m_ and the overall model goodness-of-fit for attended speech. This is not surprising, as higher SSQ_m_ scores reflect greater subjective difficulty with SIN perception, which would be expected to be related to poorer tracking of attended speech in the presence of competing speech. However, we found no correlation between SSQ_m_ and performance on the comprehension task (r = −0.17, p = 0.29), suggesting that even participants with potentially more deteriorated representations of attended speech had sufficient fidelity of speech representations to achieve high task performance. The lack of a relationship between subjective SIN perception difficulties and performance is unintuitive, but mirrors similar results showing only a weak relationship between subjective and objective measures of SIN difficulties (Phatak et al., 2018; Smith et al., 2019).

While statistical associations between subjective and objective measures of speech perception have generally been poor in past work, it is possible that these outcomes are a result of insufficiently sensitive methods for measuring speech perception. Specifically, typically used methods for objectively measuring speech perception involve presentations of isolated sentences, and having participants repeat them back, usually without time constraints (i.e., allowing participants to deliberate and piece together their percept). While these measures are simple and effective in measuring speech perception deficits in populations with moderate and severe hearing loss (e.g., Phatak et al., 2018), the external validity of these measures may be limited at best, as they do not reflect real-world listening scenarios. Specifically, real-world spoken communication generally requires real-time comprehension of complex, multi-sentence expressions embedded in noisy and reverberant backgrounds, in order to allow for continuous flow of interaction. Unlike the commonly used speech understanding tasks, these realistic scenarios allow little time for deliberation about individual words, as new information is continuous, creating the possibility of falling behind if speech processing is impaired or slowed. Indeed, Xia et al. (2017) demonstrated marked differences in performance between tasks involving simple word identification and answering comprehension questions about the content of narrative stories, with the latter showing a weaker benefit from hearing aids. This highlights the possibility that traditional speech recognition tasks may indeed be missing important, behaviorally relevant aspects of speech perception.

In the present study, a continuous multi-talker design with a behavioral task focused on assessing comprehension was selected in an attempt to mimic some aspects of real-world speech perception scenarios. Nevertheless, there were important differences that may have contributed to our failure to detect a relationship between subjective SIN perception difficulty (reflected in SSQ_m_) and behavioral performance. First, although we utilized co-located target and distractor speakers, which are generally more challenging to parse out than spatially-separated speakers (Marrone et al., 2008; Kidd et al., 2010), their fixed location, predictable temporal characteristics (e.g., lack of sudden offsets and onsets in speaking), and relatively monotone speaking styles likely facilitated participants’ ability to suppress unwanted processing of the ignored speaker. In contrast, realistic conversational settings such as restaurants or bars generally contain distractor signals that vary less predictably in location, intensity, emotional content, and other characteristics, likely contributing to greater distraction and informational masking. It is possible that suppression of these types of distractor information becomes impaired with age due to deterioration of attentional and other cognitive resources. Second, although we attempted to quantify comprehension, as opposed to mere word identification, of the content spoken by the target speaker via multiple-choice questions, it is possible that the implementation of this task lacked sensitivity to detect speech comprehension deficits. Specifically, the fact that the target story spanned many minutes may have allowed the participants to utilize much longer semantic context to aid the interpretation of incoming information, compared to real-world interactions where topics often change more rapidly. This was compounded by the fact that, for practical purposes, the questions were framed in a Yes/No format, only requiring participants to identify the more likely of the two options, rather than to demonstrate their own understanding of the story. While the main purpose of the comprehension questions was to verify that participants followed the task instructions, future work should take steps towards optimizing behavioral measures of comprehension. For example, questions carefully calibrated to require roughly constant reading time could be used to measure reaction times in addition to mere percent correct measures, possibly revealing significant response delays in people with self-reported SIN difficulties.

### 4.4 Limitations

Although our work provides evidence of age-related differences in cortical tracking of word-level features, a notable limitation of our method is that it does not establish the source of this difference. Specifically, it is unclear from our data if the distinct patterns of feature-tracking were a result of higher-order linguistic mechanisms receiving inputs with differing fidelities from lower-level processes, or they reflected age-related changes in the higher-order mechanisms themselves, or some combination of the two. Furthermore, differential engagement in cognitive resources (e.g., due to differential effort) may also have contributed to the observed differences, even in the absence of actual changes in the underlying mechanisms. Thus, an important goal for future work is to characterize speech representations more thoroughly at multiple levels of the processing hierarchy in order to elucidate the mechanisms implicated in the differences in speech processing. Furthermore, the measurement of speech representations at multiple stages of the language processing hierarchy may be critical for explaining individual differences in speech perception performance, and subjective measures such as the SSQ_m_.

The use of artificial neural networks (ANNs) to extract abstract features related to lexical and semantic content of speech has become increasingly common in studies of language processing (Huth et al., 2016; Broderick et al., 2018; Weissbart et al., 2019; Donhauser and Baillet, 2020). While powerful in characterizing brain responses to speech, an important limitation in the use of these features is that it can be difficult to interpret what aspects of language they actually capture. Specifically, ANNs are usually trained on a task such as text prediction on the basis of preceding context, and as such, ANNs may utilize any number of statistical regularities in the training corpus in order to optimize their performance. Thus, depending on the ANN architecture, aspects of language including the syntactic structure, lexical frequency, semantic relationships, and others may all contribute to the performance of ANNs. Without knowing the language aspects learned by ANNs, it is difficult, and may be even impossible, to parse out the relative contributions of the different variables. Consequently, when cortical responses are found to track these features, as is the case in the present study, it may remain unclear what linguistic processes underlie this tracking. Thus, improving the interpretability of neural analyses that utilize complex natural language models remains an important challenge for future work.

## 5 Conclusions

The present study extends upon the existing body of work demonstrating the plausibility of measuring cortical tracking of high-level features related to speech meaning and predictability. The results show evidence of age-related amplification in tracking of these features in competing speech streams. Moreover, our exploratory analyses showed trends of correlations between these measures and behavioral measures including comprehension performance and subjective SIN perception difficulty scores, indicating their potential behavioral relevance. Taken together, our work demonstrates the utility of modeling cortical responses to multi-talker speech using complex, word-level features and the potential for their use to study changes in speech processing due to aging and hearing loss.

## Data availability

Data is not available publicly, as data sharing was not a part of the informed consent. Requests to access the dataset should be directed to JM.

## Ethics statement

The Institutional Review Board of the University of Minnesota approved the procedures in this study. All participants provided written informed consent to participate.

## Author contributions

JM and MW designed the experiment, analyzed the data, and wrote the manuscript. JM and LAR implemented experimental procedures and collected the data. All authors commented on the manuscript and approved the submitted version.

## Funding

Financial support for this work was provided by NIH grant R01 DC015462 to MW. The work of LAR was supported by the NSF REU grant 1757390.

## Acknowledgments

We thank Brian Simensen and Franciska Hauer for help with data collection, PuiYii Goh for help with data analysis, Hao Lu for help with initial setup of NLP libraries, Erin O’Neill for help with designing the modified SSQ questionnaire, and Emily Allen for help with graphic design and post-processing of figures.

## Conflict of Interest Statement

The authors declare no conflicts of interest.

## Notes

### Competing Interest Statement

The authors have declared no competing interest.

